# Is Hsp110 boosting the basal disaggregation activity Hsp70 by enhanced entropic pulling strokes?

**DOI:** 10.1101/2025.04.01.646638

**Authors:** Mathieu E. Rebeaud, Bruno Fauvet, Paolo De Los Rios, Pierre Goloubinoff

## Abstract

Hsp70s use energy from ATP hydrolysis to unfold protein structures and solubilize stable aggregates, accumulating native species even under adverse non-native conditions. To carry out its catalytic polypeptide-unfolding activity, Hsp70 needs to reversibly interact with a J-domain (JDP) catalyst, a misfolded or alternatively folded polypeptide substrate and a Nucleotide Exchange Factor (NEF), which binds to the Nucleotide Binding Domain (NBD) of HSp70, accelerates ADP-release and allosterically controls the dissociation of the unfolded polypeptide product of the unfolding reaction. Yet, during the process of eukaryotisation, GrpE was lost from the cytosol, to be replaced by novel NEF proteins, among which the Hsp110 family stands out. Hsp110s belong to the Hsp70 superfamily, but the evolutionary steps that led from an ancestral Hsp70 unfoldase to a Hsp110 NEF of Hsp70s remain unsolved. Combining experiments using wild-type Sse1 (yeast Hsp110) and rationally designed mutants, we show that Hsp110 is likely built upon some of distinctive features already present in Hsp70 by repurposing them, rather than by inventing novel molecular properties. Taking all results together, we suggest a novel mechanism of action of Hsp110, whereby it is a NEF that also enhances the unfolding/disaggregating entropic pulling forces generated by Hsp70, by transiently increasing the chaperone effective volume.

## Introduction

Genome analysis of simple and more complex free-living bacteria and archaea suggests that about 3 billion years ago, early terrestrial bacteria evolved the first Hsp70 chaperone from a preexisting actin/sugar hexokinase gene (Bork, Sander et al. 1992). Apart from very simple Hsp70-less archaea and bacteria, today, Hsp70s (DnaK in prokaryotes) are present in all the ATP-containing cellular compartments of cells. Members of the Hsp70 family may accumulate into being 1% of the total protein mass of eukaryotic cells (Fauvet, Finka et al. 2021). It acts as a polypeptide unfolding nanomachine that uses energy from ATP-hydrolysis to convert stable alternatively-folded, or misfolded polypeptides, into transiently unfolded intermediates, which upon chaperone dissociation, may spontaneously reach different conformations, such as the native state, even under non-native conditions favoring misfolding and aggregation (De Los Rios and Barducci 2014, Goloubinoff, Sassi et al. 2018, Imamoglu, Balchin et al. 2020, Tiwari, Fauvet et al. 2023). Hsp70 co-evolved with obligate sub-stoichiometric (5-10 fold less) J-domain proteins (JDPs, sometimes also generically called DnaJs or Hsp40s), which accelerates ATP-hydrolysis and catalyze Hsp70’s binding to misfolded or alternatively-folded polypeptides in need to be transiently unfolded, and with GrpE that catalyzes ADP release (and ATP rebinding) and the consequent opening of the protein binding lid of Hsp70, leading to the dissociation and native refolding of the polypeptide (Russell, Wali Karzai et al. 1999, Kityk, Kopp et al. 2018). In complex bacteria and archaea, the DnaK-DnaJ-GrpE (KJE) system soon became the central hub of cellular protein homeostasis and, about a billion years later, it evolved and diversified in the cytosol and the *endoplasmic reticulum* (ER) of the first eukaryotes. Today, the Hsp70-JDP system regulates various key cellular functions necessitating structural shifts and changes of oligomeric states, affecting the biological activities of various polypeptides (Mayer and Bukau 2005, Rosenzweig, Nillegoda et al. 2019).

A simple phylogenetic analysis reveals that early in the process of eukaryotisation, an ancestral archaeal DnaK, which turned into being the first cytosolic Hsp70, did so while renouncing its billion-years dedicated nucleotide exchange factor (NEF) GrpE, for various new NEFs, such as armadillo (Fes1 in yeast) and Bag1 (Snl1 in yeast)(Bracher and Verghese 2015), for which no obvious orthologous prokaryotic ancestor can be identified (Rebeaud, Mallik et al. 2021). Like GrpE, both NEFs don’t hydrolyze ATP. Upon transiently binding to Hsp70’s nucleotide binding domain (NBD), both increase ADP-release and ATP binding and promote the release of the Hsp70-bound polypeptides (Rosenzweig, Nillegoda et al. 2019). Importantly, the first eukaryotes (Richards, Eme et al. 2024) also evolved a new NEF, called Hsp110, from a prokaryotic Hsp70 ancestor (DnaK), which became the most abundant NEF in the yeast and human cytosols. Soon after the appearance of the first cytosolic Hsp110, an orthologous version of it further evolved by gene duplication, to be exported and serve in the *endoplasmic reticulum* (ER) as the sole NEF (called Lhs1 in yeast, Hyou1 or Grp170 in mammals) to the newly ER-exported Hsp70s (called BIPs) (Rebeaud, Mallik et al. 2021). About two billion years later, all Hsp110s in current eukaryotes are highly homologous, sequence-wise and structurally, to their Hsp70 ancestor.

Here, we aimed to identify the transformations that changed an original Hsp70 chaperone, which was an ATP-fueled JDP-dependent polypeptide-unfolding enzyme that binds NEFs, into Hsp110, which is an ATP-fueled JDP-independent nucleotide exchange catalyst of Hsp70, devoid of polypeptide-unfolding abilities by itself and that does not bind NEFs. Our approach is to use wild-type (WT) yeast SSE1 and some designed mutants (Sup Fig. 2 and 3), and measure the ATPase, prevention of aggregation, unfolding, disaggregation and refolding activities of Ssa1, Sis1 and Sse1, respectively the most abundant Hsp70, JDP and NEF cochaperones in the cytosol of yeast cells. We find that the Hsp110s have apparently renounced their ancestral ability to interact with, and have their ATPase activated by J-domain cochaperones (JDPs)(Goeckeler, Petruso et al. 2008, Kumar, Peter et al. 2020). Without Hsp70s, Hsp110s have also apparently forsaken most of their ability to bind misfolded polypeptides, interact with NEFs and to remodel various polypeptide substrates by ATP-fueled unfolding. Yet, the Hsp110s have apparently gained the ability to form transient dimers (Shaner, Sousa et al. 2006, Liu and Hendrickson 2007) with Hsp70s through their conserved nucleotide binding domain (NBD) and, at variance with GrpE, oddly depend on ATP hydrolysis to accelerate ADP release from Hsp70 (Raviol, Bukau et al. 2006).

## Results

### Phylogenetic analysis of Hsp110s

Ssa1 and Sse1 are the most abundant Hsp70 and Hsp110 molecules in the cytosol of exponentially growing yeast, estimated to be 16 µM and 3 µM, respectively (Ghaemmaghami, Huh et al. 2003, Brownridge, Lawless et al. 2013, Lawless, Holman et al. 2016, Mackenzie, Lawless et al. 2016) (Supplementary Table 1). Ssa1 has three orthologues, Ssa2, Ssa3 and Ssa4, which together account for less than two micromolar. Likewise, the Hsp110 orthologue, Sse2, is less than 0.6 µM (Ghaemmaghami, Huh et al. 2003, Brownridge, Lawless et al. 2013, Lawless, Holman et al. 2016, Mackenzie, Lawless et al. 2016). ADP-bound Hsp70 and ATP-bound Hsp110 can form heterodimeric crystals (Shaner, Sousa et al. 2006, Polier, Dragovic et al. 2008). AlphaFold 3 predictions (Jumper, Evans et al. 2021, Manalastas-Cantos, Adoni et al. 2024) confirmed this nucleotide preference for the Ssa1-Sse1 heterodimer (Fig. 1 A). The observed natural 5-7-fold excess of Ssa1 over Sse1 in the yeast cytosol suggests that, whereas most Sse1-2 can be found bound to Hsp70, at least 80% of the Hsp70s must be free of Hsp110 at any moment. This requires heterodimers to be dynamic and to drive an optimal chaperone activity that likely depends on multiple cycles of Sse1 binding to and dissociating from Ssa1.

**Figure 1:**
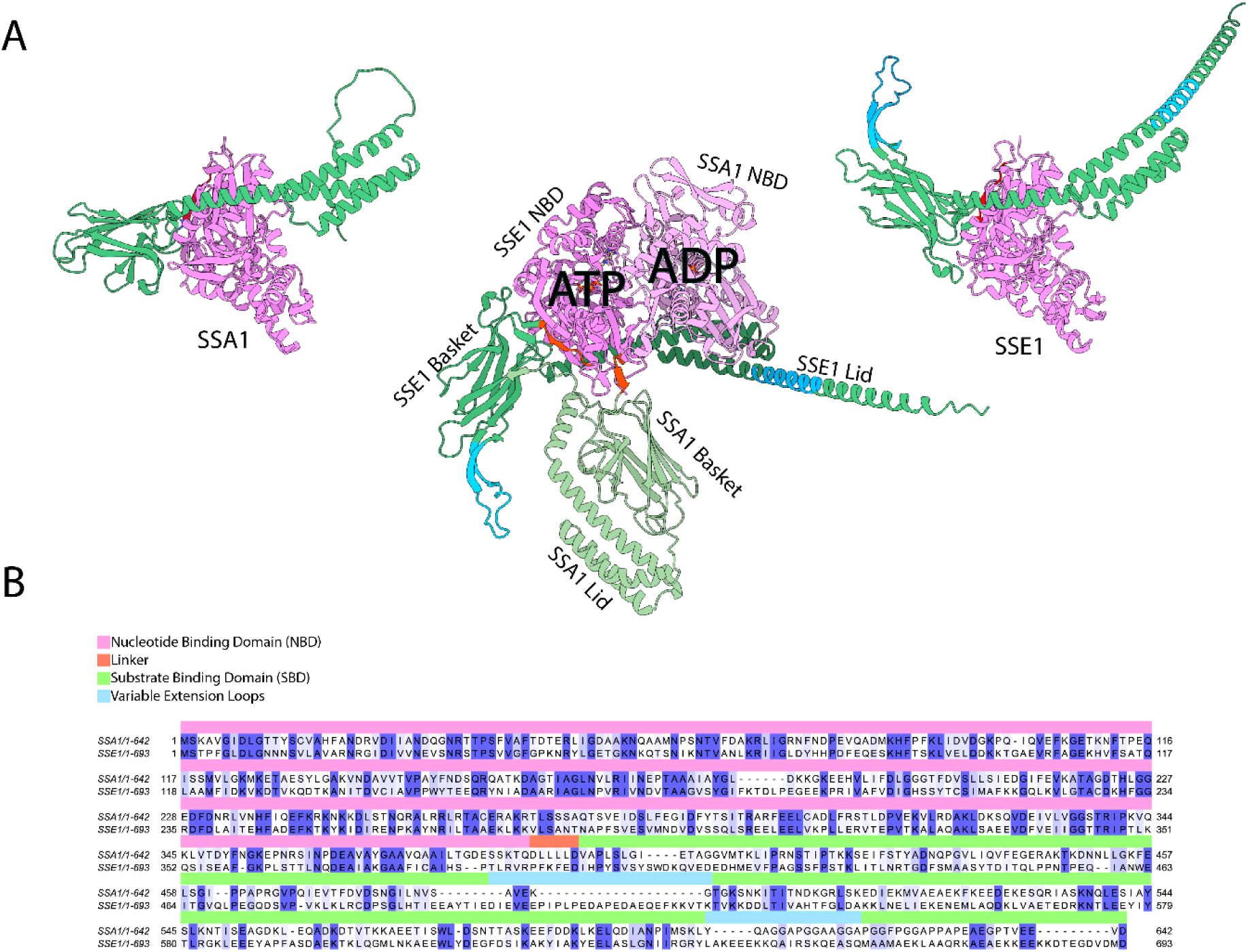
Structures and sequences of yeast Ssa1 and Sse1. **A**: AlphaFold and crystal structures (UNIPROT: AF-P32589-F1 and AF-P10591-F1) of ATP-bound SSA1 (left) and ATP-bound SSE1 (right) and their hetero-dimeric complex in the middle. AlphaFold top suggestion for an Ssa1-Sse1 heterodimer places ATP in Sse1 and ADP in a “relaxed” conformation of Ssa1, with a closed SBD that can potentially pull misfolded polypeptides out of a stable aggregate. Ssa1 and Sse1 are associated though their Nucleotide Binding Domain (NBD, Magenta). Ssa1’s Substrate Binding Domain (SBD, Green) is composed of a β-folded “basket” subdomain and an α-helical lid that can close onto a misfolded substrate. Non-homologous extended segments in Sse1’s basket and lid are in pale blue. **B**: Amino acid sequence alignment of yeast SSA1 and SSE1 showing the conserved NBDs (violet), distinct likers (red) and highly variable extra-loops (blue) in the less conserved SBD (green). Amino acid conservation scores from 50 to 100% are highlighted in a blue gradient.

**Supplementary Figure 1:**
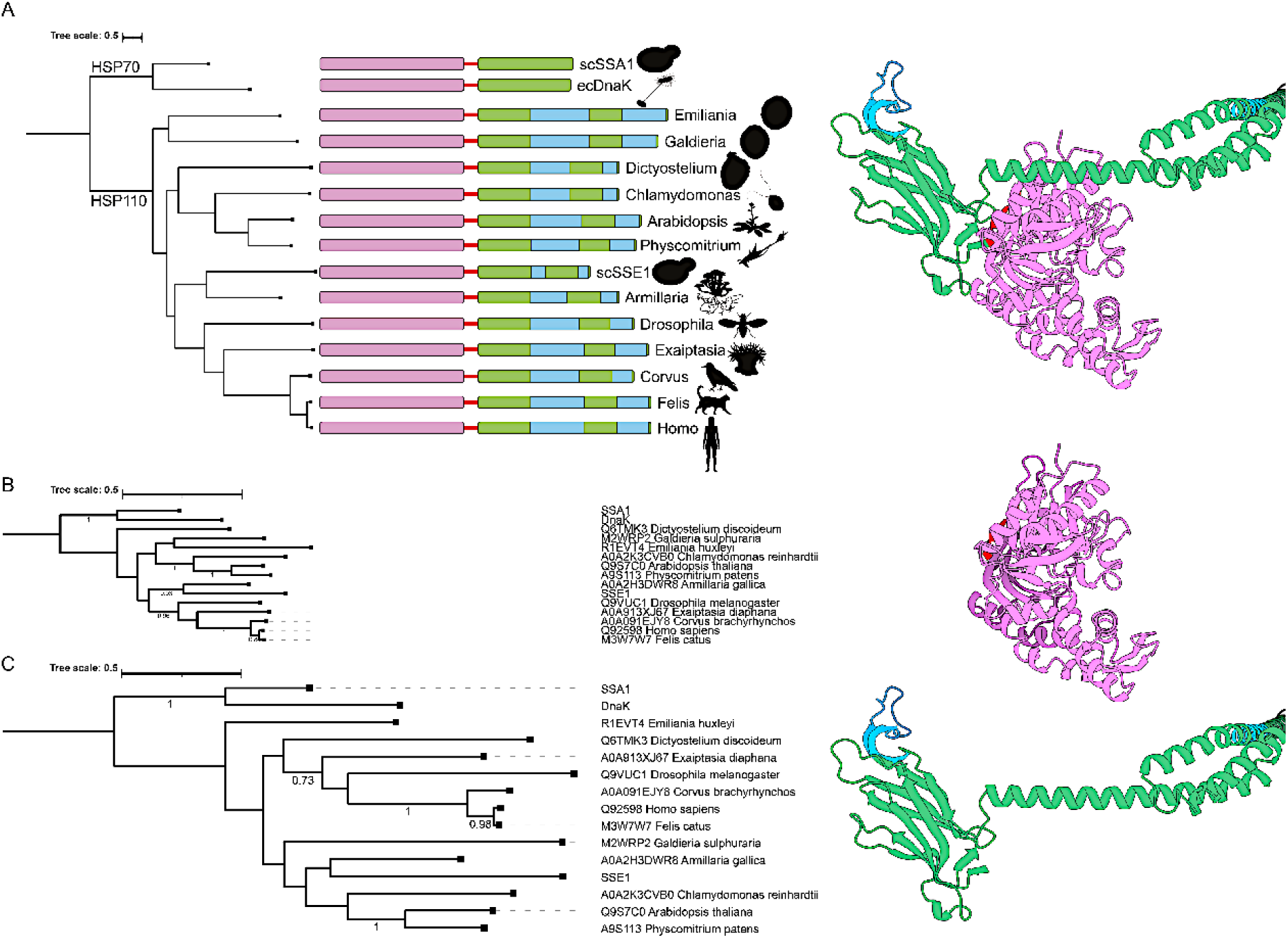
Phylogenetic trees from amino acid sequence alignments of HSP110s from distant eukaryotes, rooted with *E. coli* DnaK and Yeast HSP70s. (A) Compared to the highly conserved HSP70s, the SBDs of HSP110’s contain increasingly long, variable extra loops (blue) in the SBD, which in yeast SSE1 are the shortest among all the known HSP110s (right scheme). Sequence variation show during the last ∼2 billion years of eukaryote evolution, amino acid substitutions were much slower in the NBDs (B, SSE1 NBD in violet), than in the SBDs (C, SSE1 SBD in green) of both Hsp70s and Hsp110s. Bootstrap value of more than 0.7 are shown on the trees of B and C (Representative organisms pictures are from https://www.phylopic.org, with *Chlamydomonas* by Sergio A. Muñoz-Gómez and *Escherichia coli* by Matt Crook)

**Supplementary Figure 2.**
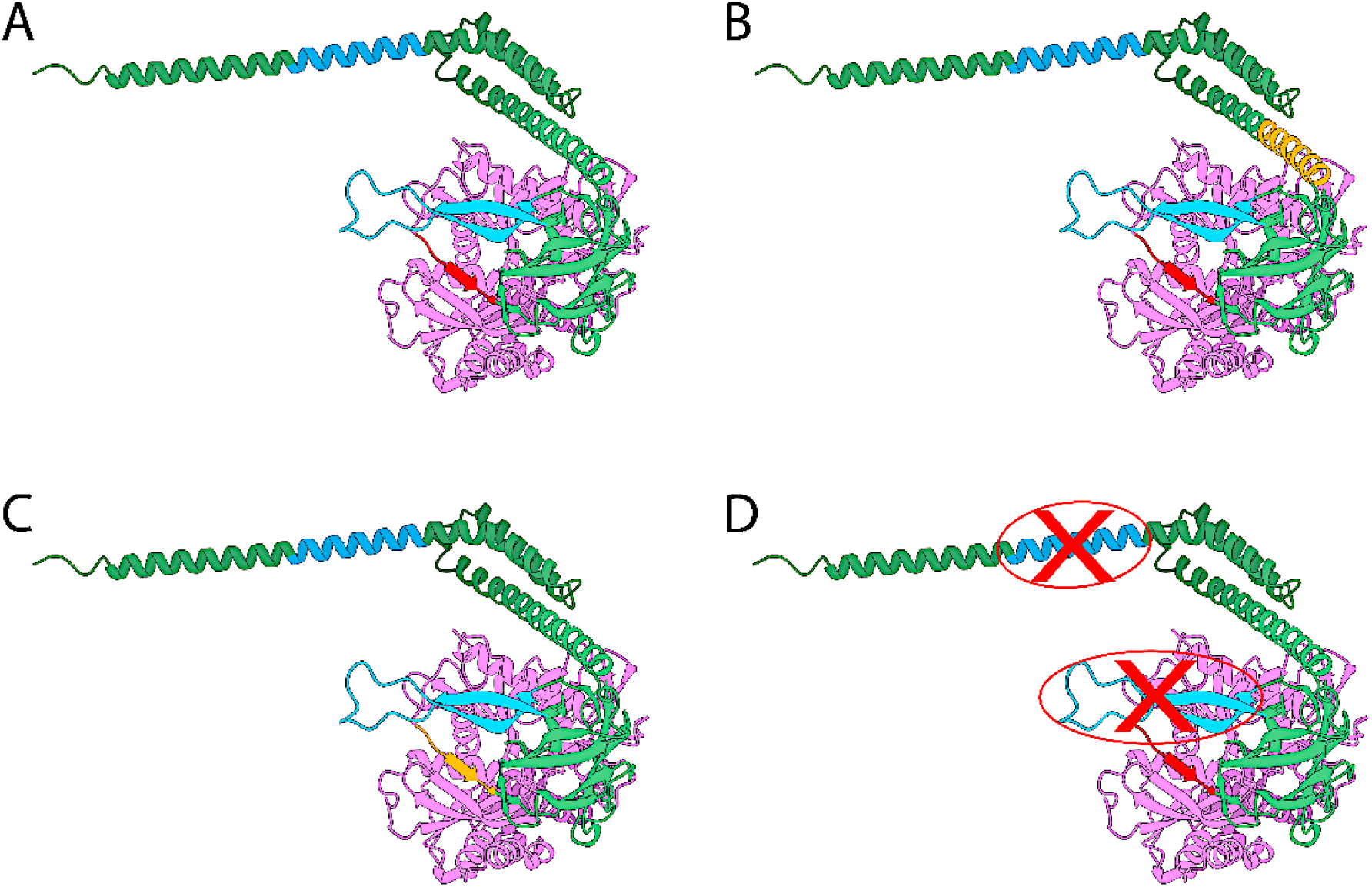
Structure of SSE1 WT with changes introduced in the mutants used in this study. **A**: WT SSE1 AlphaFold-generated structure (from UNIPROT: AF-P32589-F1), with NBD in purple, linker in red, SBD in green and the extensions in blue. **B**: SSE1 HM (helix mutant, orange). **C**: SSE1 LM (linker mutant, EAE, orange). **D**: SSE1 Loopout, with the extension crossed in red.

**Supplementary Figure 3.**
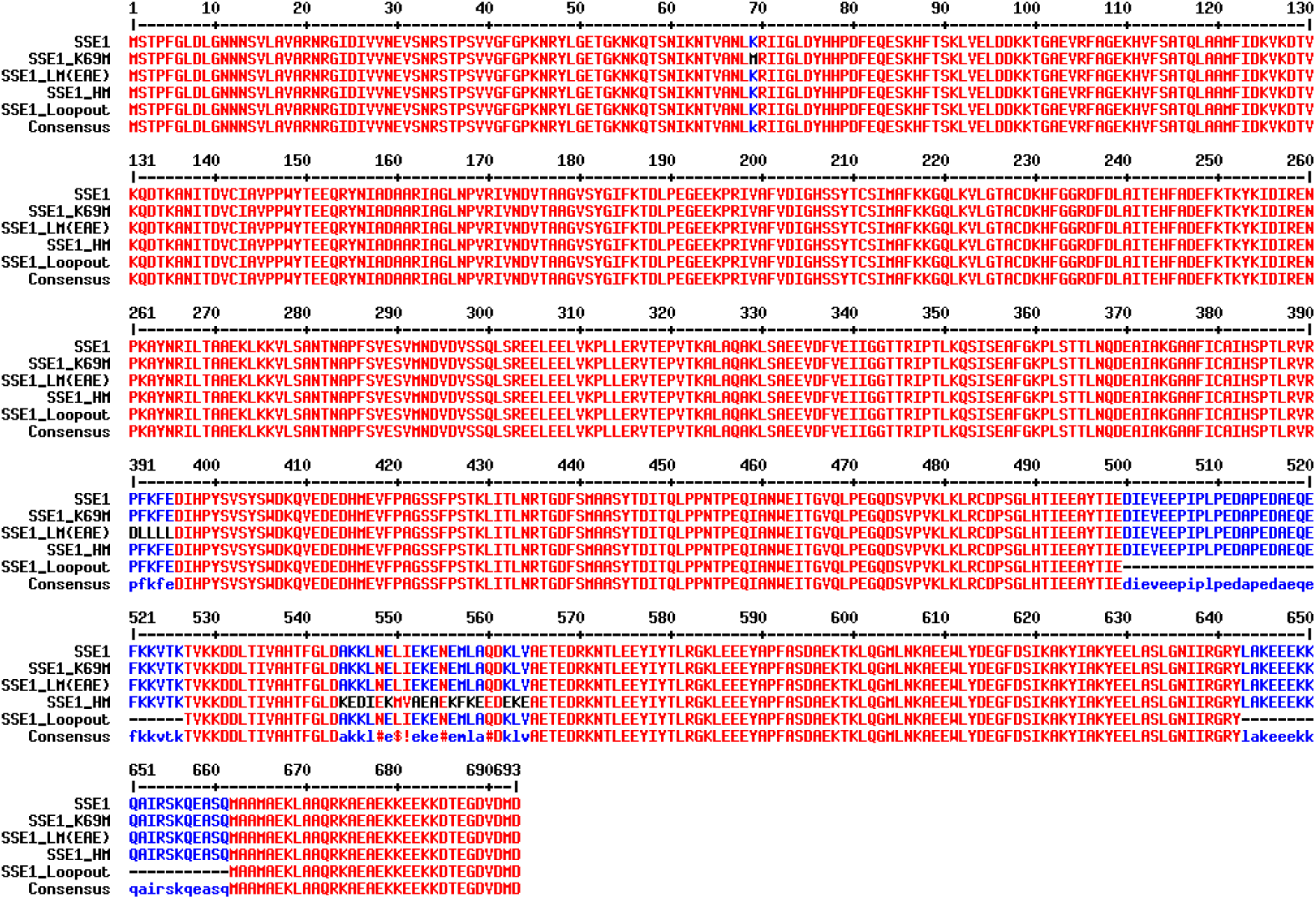
Sequence alignment of SSE1 with the mutants used in this study. Red, identical residues. Black, residues from Ssa1. Blue, identical residues missing in the Loopout mutant or different in the EAE and HM mutants.

Sequence-wise, Ssa1 and Sse1 are highly homologous. Yet, Sse1 presents a distinctive linker between its NBD and SBD domains (PFKFED), which is a hallmark for yeast Hsp110s (Chakafana, Mudau et al. 2021). This linker is clearly distinguishable from the highly conserved linker sequence of prokaryotic and eukaryotic Hsp70s, composed of a core of four small hydrophobic, non-aromatic residues flanked by negative charged residues (DLLLLD for Ssa1) (Liu and Hendrickson 2007, Chakafana, Zininga et al. 2019) (Fig. 1B). A phylogenetic tree of amino acid sequence alignment of Hsp110s from distant eukaryotes, using *E. coli* DnaK and Yeast Ssa1 as outgroups, reproduced the known structure of the tree of life (Sup Fig. 1C). The phylogenetic trees for the two separate domains confirmed that Hsp110’s NBDs are much more conserved than their SBDs (Sup Fig. 1). There are two highly variable regions in Hsp110’s SBD, one in the so-called β-strand “basket”, and another in the α-helical “lid”, with unstructured extended segments, according to AlphaFold (Sup Fig. 1). To simplify our search for what in general tells Hsp110s apart from Hsp70s, we chose Sse1 as a model Hsp110 because it has the shortest extensions in the SBD, while retaining all the other features of the Hsp110s family.

### Sse1 by itself cannot prevent MlucV aggregation

Molecular chaperones are often thought to act as “holdases”, by way of binding stress-unfolded polypeptides, thereby preventing their further aggregation (Sontag, Samant et al. 2017, Rosenzweig, Nillegoda et al. 2019). We monitored the individual and combined abilities of the main yeast cytosolic Hsp70 (Ssa1), JDPs (Sis1 and Ydj1) and NEF (Sse1), to prevent the aggregation of urea pre-unfolded MlucV (Tiwari, Fauvet et al. 2023). This reporter protein was previously developed to assess the native/non-native state of thermolabile mutated firefly luciferase (Luc)(Sharma, De Los Rios et al. 2011) through its enzymatic activity, and its compactness by Förster Resonance Energy Transfer (FRET) between the N-terminal (M = mTFP1) and C-terminal (V = Venus) fluorescent proteins (Tiwari, Fauvet et al. 2023).

The luciferase core of MlucV was first denatured in 4.8 M urea. Then, urea was extensively diluted by adding buffer and ATP in the absence or presence of increasing amounts of Ssa1, without or with Sis1 and/or Ydj1 and/or Sse1 (Fig. 2A). After dilution, thus in conditions favoring the stability of the native state, in the absence of chaperones the luciferase core of MlucV was found to be, as expected, enzymatically inactive, while the FRET values from the flanking urea-resistant native GFP-type fluorophores were typical of stable soluble MlucV aggregates, namely with about 150% of the FRET level set at 100% for the native MlucV)(see (Tiwari, Fauvet et al. 2023)). When dilution and aggregation occurred in the presence of increasing amounts of Ydj1, a class A JDP similar to *E. coli* DnaJ (Kampinga, Andreasson et al. 2019), the FRET signal was gradually lowered from 150% without Ydj1, to ∼105% at the largest Ydj1 concentrations. This is owing to the well-established ability of Ydj1 to bind polypeptides that misfold and thereby prevent their further aggregation (Lu and Cyr 1998, Rebeaud, Tiwari et al. 2024). In contrast, up to a 24 folds molar excess of Sis1, a class B JDP similar to *E. coli* CbpA (Kampinga, Andreasson et al. 2019), left the high FRET signal essentially unaltered, consistent with Sis1’s recognized inability to strongly bind unfolded/misfolded monomers and prevent their aggregation (Rebeaud, Tiwari et al. 2024). Like Sis1, Sse1 proved to be a very poor “holdase” by itself. Less expectedly, increasing amount of Ssa1, the canonical and most abundant cytosolic yeast Hsp70, only very mildly reduced the FRET signal of MlucV, down to ∼145% at the largest measured concentrations. In none of these cases the individual action of the chaperones and cochaperones resulted in a detectable recovery of the enzymatic luciferase activity, despite the presence of ATP in the assay (Fig. 2B), suggesting that the mild FRET reduction was associated with a very minor prevention of aggregation that did not subsequently lead to any refolded native species.

**Figure 2:**
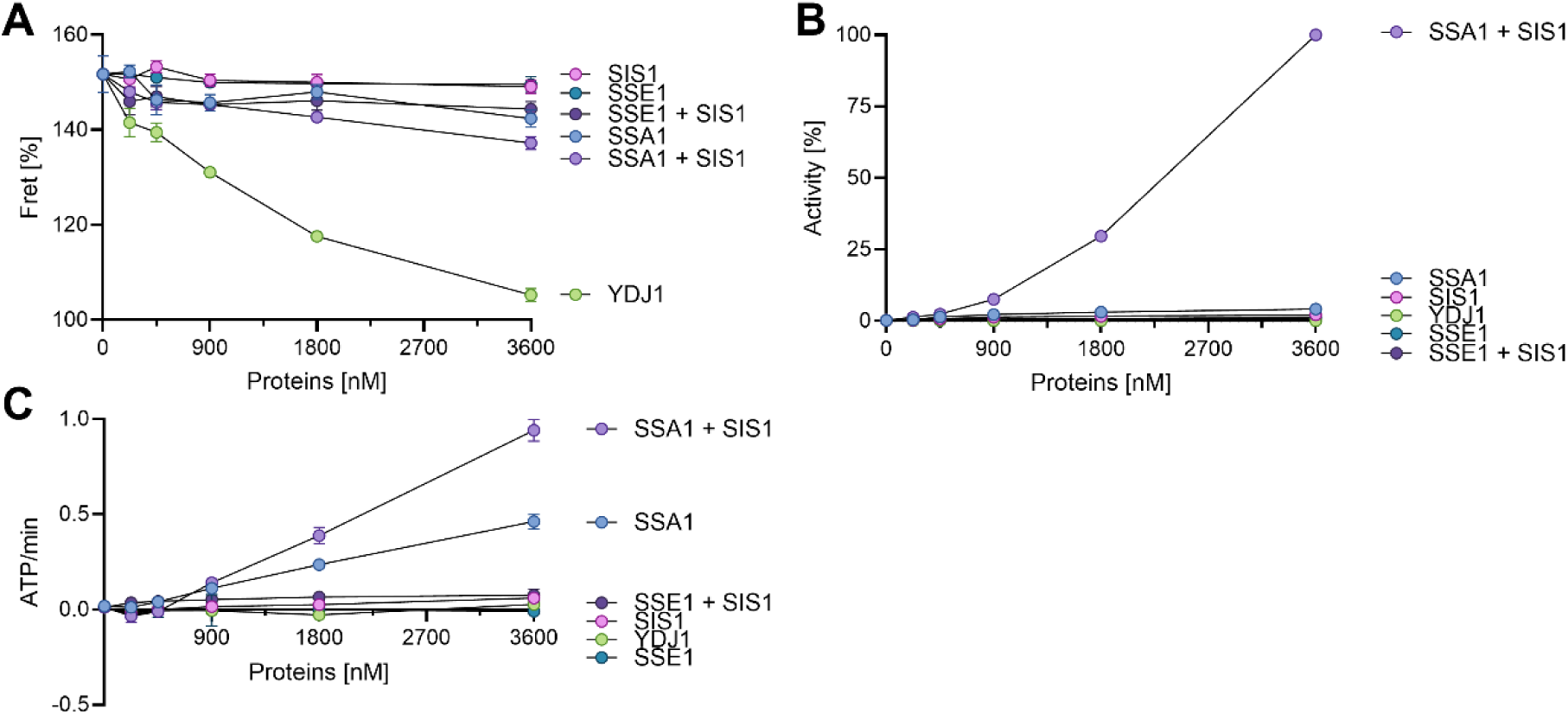
The effect of increasing amounts of SSA1 or Sse1, with MlucV aggregates, without or with Sis1 on ATP hydrolysis, the aggregation and the conversion of aggregates into native MlucV. Urea-pre-unfolded MlucV was diluted in the presence of ATP and increasing concentrations of either Ssa1, or Sse1, without or with 900 nM Sis1.Increasing Ydj1 was also measured as a control. **A**: The FRET values of urea-pre-unfolded MlucV diluted in the presence ATP and increasing concentration of Ydj1, or Ssa1 or Sse1, without or with 900 nM Sis1. **B**: Regain of native luciferase from urea-pre-unfolded MlucV, following 80 min in the presence of the different chaperone and co-chaperone mixes as in A. Maximum relative luciferase reactivation was set at 100%. **C**: ATPase activity of the different chaperones mixes as in A.

We next tested the effects of increasing amounts of Ssa1 or Sse1 in the presence of constant (900 nM) Sis1 (that by itself, virtually does not prevent aggregation), finding that their effects on urea-pre-unfolded, misfolding MlucV were very different: Whereas both chaperone combinations reduced the FRET signal slightly more than the individual proteins (Fig. 2A), there was no recovery of luciferase activity by Sse1+Sis1 at any concentration while, in contrast, increasing concentrations of Ssa1 with Sis1 led to a mild but significant reduction of the FRET signal and a corresponding increment of the recovered luciferase activity (Fig. 2C). We also measured the ATPase activity of increasing concentrations of Sse1 or Ssa1, with aggregated MlucV, without and with Sis1 (Fig. 2B). Not surprisingly, the number of hydrolyzed ATP molecules per minute was undistinguishable from the background for Sis1 and Ydj1 alone, since they are not ATPases. Sse1, alone or in combination with Sis1, did not hydrolyze detectable amounts of ATP, despite having retained an intrinsic ATPase activity (Kumar, Peter et al. 2020). By converse, increasing concentrations of Ssa1, showed a detectable ATPase activity, which doubled in the presence of Sis1.

### Without NEFs, Hsp70s-JDPs, have a significant disaggregase/refolding activity

In the presence of Sis1, Ssa1 weakly but effectively binds pre-aggregated polypeptides and uses energy from ATP-hydrolysis to dissolve them by unfolding through entropic pulling forces (De Los Rios, Ben-Zvi et al. 2006, Wyszkowski, Janta et al. 2021, Tiwari, Fauvet et al. 2023). According to this model, a NEF is expected to merely accelerate ADP dissociation and ATP rebinding, cause a consequent opening of Hsp70’s SBD and allow the dissociation of the locally unfolded polypeptide intermediate, which can freely locally refold in solution, ultimately to the native state (Sharma, De los Rios et al. 2010, Mayer and Gierasch 2019, Imamoglu, Balchin et al. 2020, Tiwari, Fauvet et al. 2023). Given the purported NEF function of Sse1, we next addressed its functional effects on the protein disaggregation and refolding activity of the corresponding Hsp70-JDP chaperone system (Ssa1-Sis1) and compared it with the bacterial Hsp70-JDP-NEF system (DnaK-DnaJ-GrpE, KJE).

While urea-pre-unfolded and preaggregated MlucV species alone did not spontaneously convert into native MlucV over time (Fig. 3), the bacterial KJ system, together with ATP, effectively converted a large fraction of the stable MlucV aggregates into native MlucV. Having GrpE present as well (KJE) further enhanced the MlucV reactivation rates and yields, albeit only marginally.

**Figure 3:**
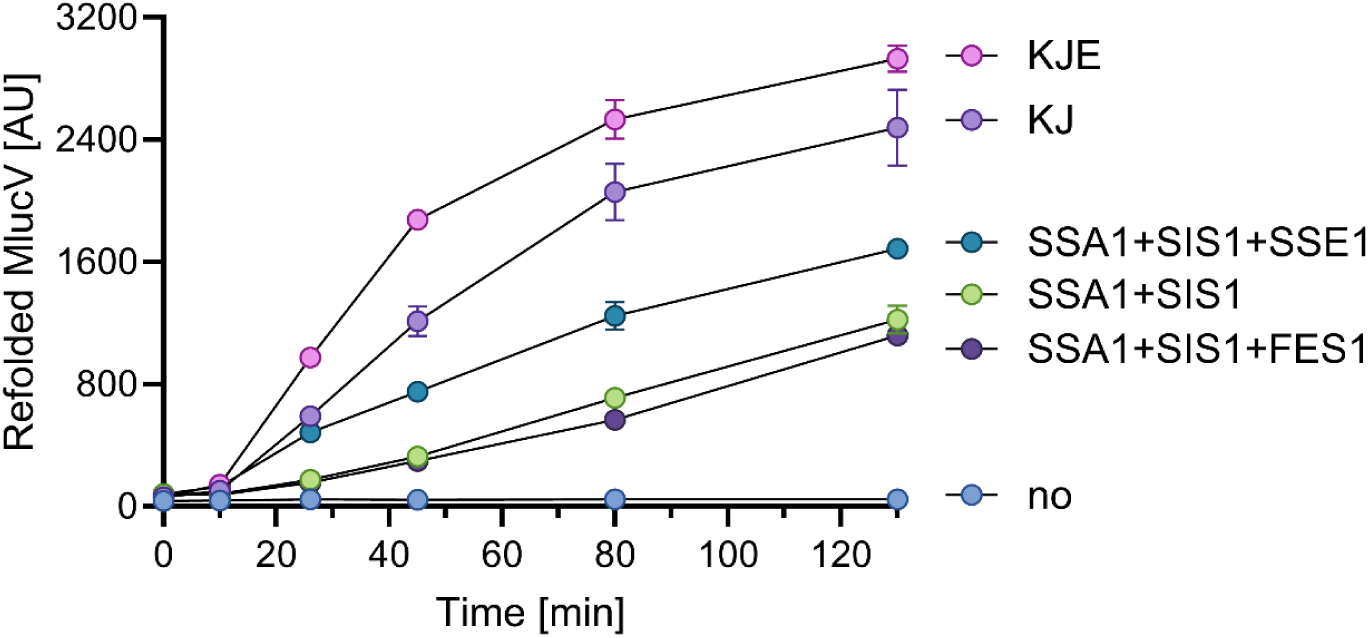
Bacterial and yeast HSP70-JDP can disaggregate and refold stable aggregates in the absence of NEFs. Time-dependent regain of luciferase activity from pre-aggregated MlucV, by yeast or *E. coli* chaperones: 150 nM of stable, inactive MlucV aggregates were incubated up to 120 min at 25 °C in the presence of 5 mM ATP, 4000 nM Hsp70 (DnaK or Ssa1), 1000 nM JDP (Sis1 or DnaJ), with or without NEF (500 nM Sse1, 1000 nM Fes1 or 1000 nM GrpE), as indicated.

Similarly, but about twice as slow as the bacterial chaperones, Ssa1 together with Sis1 but without Sse1 also effectively converted a significant fraction of the stable aggregates into native MlucV. Addition of an optimal concentration of Sse1 (500 nM, in the same Ssa1/Sse1ratio as the yeast cytosol) slightly but significantly increased the luciferase refolding rates and yields. Addition of Fes1, another NEF in the yeast cytosol, did not stimulate the intrinsic disaggregase/refolding activity by Ssa1+Sis1 alone, an effect that at the tested concentration was thus specific to Sse1. Contrary to the obligate need for a JDP co-chaperone, without which Hsp70 had no refolding activity, both prokaryotic GrpE and eukaryotic Sse1 thus appear, *in vitro*, to be less obligate cochaperones: their action accelerated the refolding rates and increased the refolding yields only to a minor extent.

We next performed a dose-response of the NEFs and measured the luciferase reactivation of pre-aggregated MlucV and luciferase after 2 hours of chaperone action. Fig.4 confirmed that both DnaK-DnaJ and Ssa1-Sis1 have a strong stand-alone disaggregation and refolding activity of their own, which does not require the presence of GrpE and Sse1, respectively. This notwithstanding, both NEFs enhanced the activity of their corresponding chaperone systems: Increasing amounts of GrpE nearly doubled the yields of the stand-alone disaggregation and refolding reaction by DnaK-DnaJ, with an EC_50_ at about 200 nM, corresponding to ∼40 times fewer GrpE dimers than DnaK monomers (Fig. 4). This is suggestive of a very dynamical mode of action, where for an optimal activation of DnaK-DnaJ a single GrpE dimer may iteratively bind, activate and dissociate from as many as 20 different DnaK molecules. Sse1 similarly showed a dynamical mode of action, since it nearly doubled the yields of the stand-alone disaggregation and refolding activity by Ssa1-Sis1, with an EC_50_ at 100 nM, when Sse1 was ∼40 times less than Ssa1 (Fig. 4). Yet, at variance with the excess of GrpE that neither further activated nor inhibited the chaperone reaction, Sse1, above 250 nM, increasingly inhibited the disaggregation and refolding reaction, with an IC_50_ at 1200 nM and a complete inhibition at the non-physiological ratio of equimolar Sse1 and Ssa1, similar to equimolar human Apg2, which was been reported to inhibit Hsc70(Cabrera, Dublang et al. 2019).

**Figure 4:**
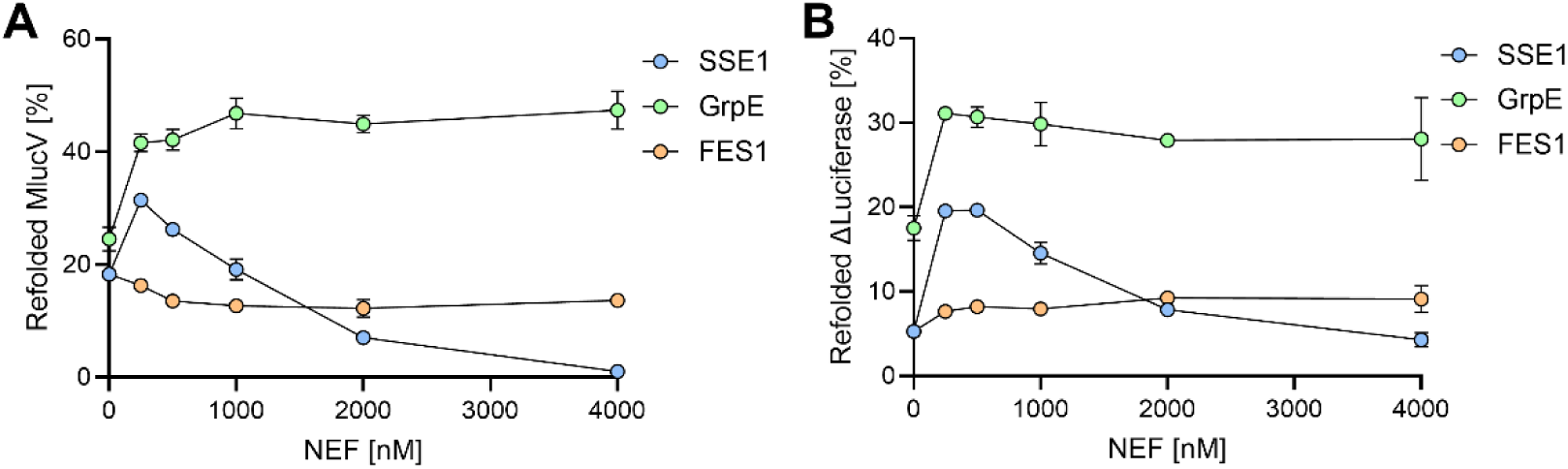
Dose-responses of bacterial and eukaryotic NEFs with different substrates. **A**: MlucV was preaggregated (150 nM) in the presence of 4 mM ATP, 4000 nM Ssa1 and 1000 nM Sis1 and increasing amounts of Sse1. For comparison, the same reaction was carried with 4000 nM DnaK,1000 nM DnaJ and increasing amounts of GrpE. **B**: Similar high yields were observed as in A but using stable inactive luciferase aggregates (without fluorophores), which form much larger and more compact insoluble oligomers than stable MlucV aggregates.

To address the process by which the stand-alone disaggregation and refolding reaction by Ssa1-Sis1 becomes inhibited by an excess of Sse1, we next measured the time-dependent changes in the FRET value and in the recovered luciferase activities of preaggregated MlucV. When incubated at 34°C with Ssa1 and Sis1 in the presence of ATP, a sharp decrease of the FRET signal of the preaggregated MlucV was observed (Fig. 5A), which confirmed that even without Sse1, the initial addition of Ssa1 and its JDP cochaperone can effectively drive the decompaction and unfolding of the stable preaggregated MlucV species and lead to the recovery of native luciferase activity (Fig. 5B). Subsequent addition at t=40 min of an Sse1 concentration close to the optimal ratio with Ssa1 (500 nM, see Fig. 4), nearly doubled the rate of MlucV refolding, and to a lesser extent the yields. As expected from the accumulation of more native MlucV, the FRET signal also slightly increased because of the conversion of low-FRET chaperone-bound unfolded species into more compact native species with a higher-FRET (Fig. 5A) (Tiwari, Fauvet et al. 2023). In contrast, the addition at t=40 min of a large, non-physiological excess of Sse1 (4000 nM, equimolar to Ssa1) abruptly increased the FRET signal (Fig. 5A), suggesting that when excessively bound by Sse1, Ssa1 was not only unable to convert bound MlucV to an unfolded conformation, but that likely it was unable to counteract the aggregation of unbound, free MlucV. Correspondingly, the luciferase reactivation of MlucV was also halted (Fig. 5B).

**Figure 5:**
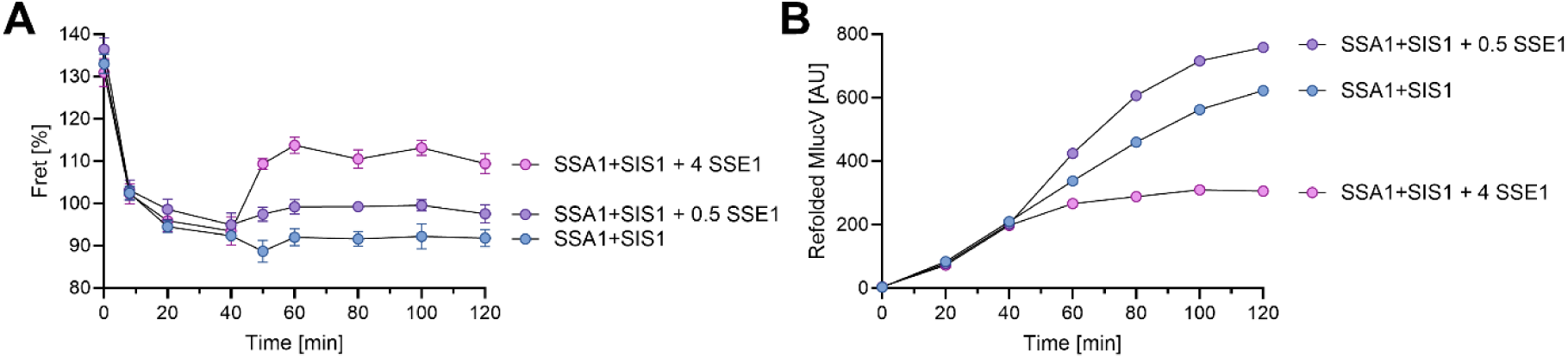
Excess SSE1 causes aggregation and prevents unfolding by SSA1-SIS1 and arrests native refolding. MlucV was pre-unfolded in urea and abruptly diluted 100-fold in refolding buffer containing 4 mM ATP and after 5 min at 25°C, 4000 nM Ssa1 and 500 nM Sis1. After 40 minutes, 500, 4000 nM or no Sse1 was added to the mix as indicated. FRET signals (A) and luciferase activity (B) were measured for 2 hours.

Similar results were obtained in the absence or presence of YS, a dysregulated hyper-active J-domain co-chaperone of Ssa1 (Rebeaud, Tiwari et al. 2024). YS is a chimeric protein comprising the J-domain of Ydj1 and the G/F linker and CTDs of Sis1. The swap of the J-domain disrupts the inhibitory interactions between the two parts, making the J-domain always available for Ssa1 binding. Like Sis1, with which it shares the CTDs, YS was unable by itself to prevent MlucV aggregation (Sup Fig. 4A). With ATP, the addition of 4000 nM Ssa1 to YS caused a decrease of the initial FRET value and a mild, yet significant luciferase refolding was measured (Sup Fig. 4B),. This indicated that most of the preaggregated substrates readily became bound to Ssa1, was prevented from further aggregating and was unfolded and subsequently released to natively refold. Expectedly, the time-dependent unfolding and native refolding was increased with the addition of an optimal, sub-stoichiometric amount of Sse1 (500 nM). Instead, and in line with the results in Fig. 5, excess equimolar Sse1 abolished the ability of the Ssa1 and YS to prevent MlucV aggregation and/or to unfold and disaggregate preformed aggregates and subsequently lead to their native refolding.

**Supplementary Figure 4:**
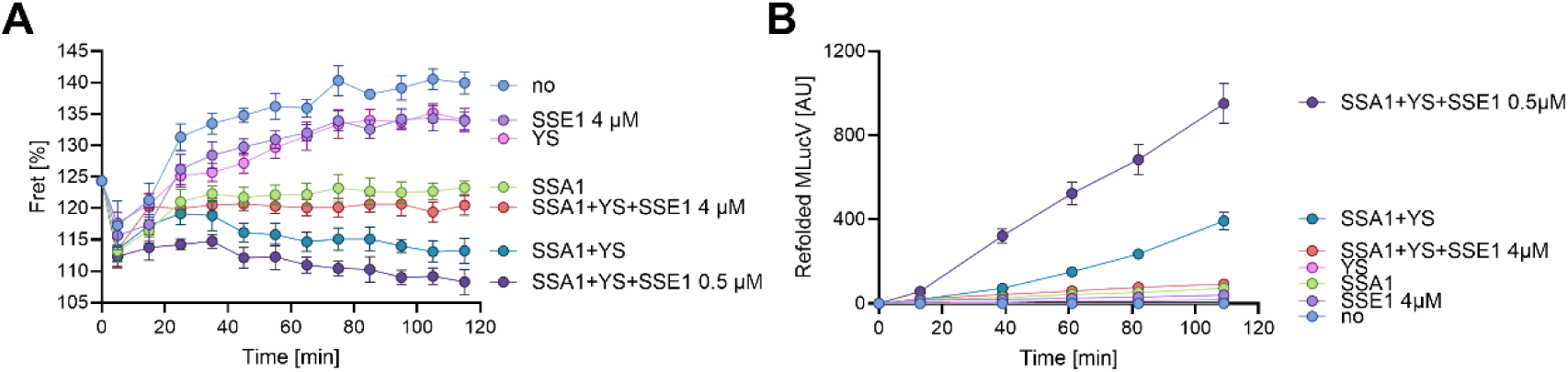
Contrary to limiting SSE1, excess Sse1 does not prevent aggregation and blocks native refolding by SSA1-SIS1. MlucV was pre-unfolded in urea and abruptly diluted 100X in refolding buffer containing 4 mM ATP and after 5 min at 25°C to allow aggregation, 4000 nM Ssa1 or 4000 nM Sse1 was added or not (no), in the absence or presence of 1000 nM Sis1, without or with 500 nM Sse1. as indicated. FRET signals (A) and luciferase activity (B) were measured for 2 hours.

### Nucleotide Exchange Factors rescue Hsp70 activity at high ADP concentrations

Despite our *in vitro* activity assays indicating that GrpE is a less essential co-chaperone than DnaJ for DnaK, in bacteria GrpE is often found associated with DnaK and DnaJ in the same operon (Genevaux, Georgopoulos et al. 2007), suggestive of its very important function. Similarly, Sse1 is constitutively expressed in the cytosol of yeast. Whereas in some extreme cases, like for the *in vitro* disassembly of compact α-synuclein fibrils, Hsp110s seem to be stringently essential (Wentink, Nillegoda et al. 2020), our results with less compact aggregates show that in general, NEFs rather play the role of optimizing the strong stand-alone disaggregation and refolding activities of Hsp70s-JDPs core-machinery. To further investigate the degree of essentiality of NEF cochaperones in bacteria and yeast, we next addressed their effect in the presence of limiting ATP and increasing ADP concentrations. Indeed, from their name, the putative function of NEFs is to facilitate the release of ADP from Hsp70 and the consequent rebinding of a new nucleotide molecule, which most likely would-be ATP in the usual ATP excess conditions, but might increasingly be ADP as its concentration increases, for example in energy depleted cells. It is important to stress that in quiescent micro-organisms, as well as under starvation, the ADP/ATP ratios can be elevated (Doello, Burkhardt et al. 2021, Takaine, Imamura et al. 2022).

There are two reasons to expect that increasing an ADP concentration relative to ATP should inhibit the Hsp70-JDP system. First, ADP locks Hsp70 in its closed state, preventing substrate release and subsequent native refolding and rebinding of a new misfolded substrate. Second, the energy available from ATP hydrolysis, which powers the chaperone machinery, is given by

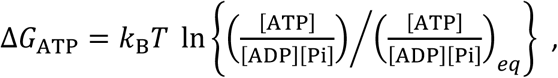

Where 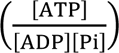 is the concentration ratio of ATP to its hydrolysis products in the chaperone assay, and 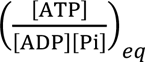 is the equilibrium ratio, if the nucleotide solution were allowed to reach thermodynamic equilibrium ([ADP]≫[ATP]). Increasing ADP reduces the first ratio, thereby depriving Hsp70 of the energy required for its function.

We thus measured the ability of either DnaK or Ssa1, in the presence of their respective JDPs and increasing amounts of ADP over a fixed near-limiting amount of ATP (0.8 mM), to natively refold stable preaggregated inactive MlucV species over time, without or with GrpE or Sse1, respectively (Fig. 7). Expectedly, in the absence of NEFs, both the prokaryotic and eukaryotic Hsp70-JDP systems were progressively inhibited by higher ADP concentrations over a constant limiting concentration of ATP (800 nM), with an IC_50_ for [ADP]/[ATP]≈2 (Fig. 6A). More surprisingly, the presence of sub-stoichiometric amounts of NEFs (3 and 7.5 times less in the case of GrpE and Sse1, respectively), significantly alleviated the inhibition by an excess of ADP over ATP. In the case of Ssa1-Sis1 with Sse1, equimolar ADP and ATP caused a slight, but reproducible, increase in the refolding yields (Fig. 6A and B) and rates (Fig. 6C).

**Figure 6:**
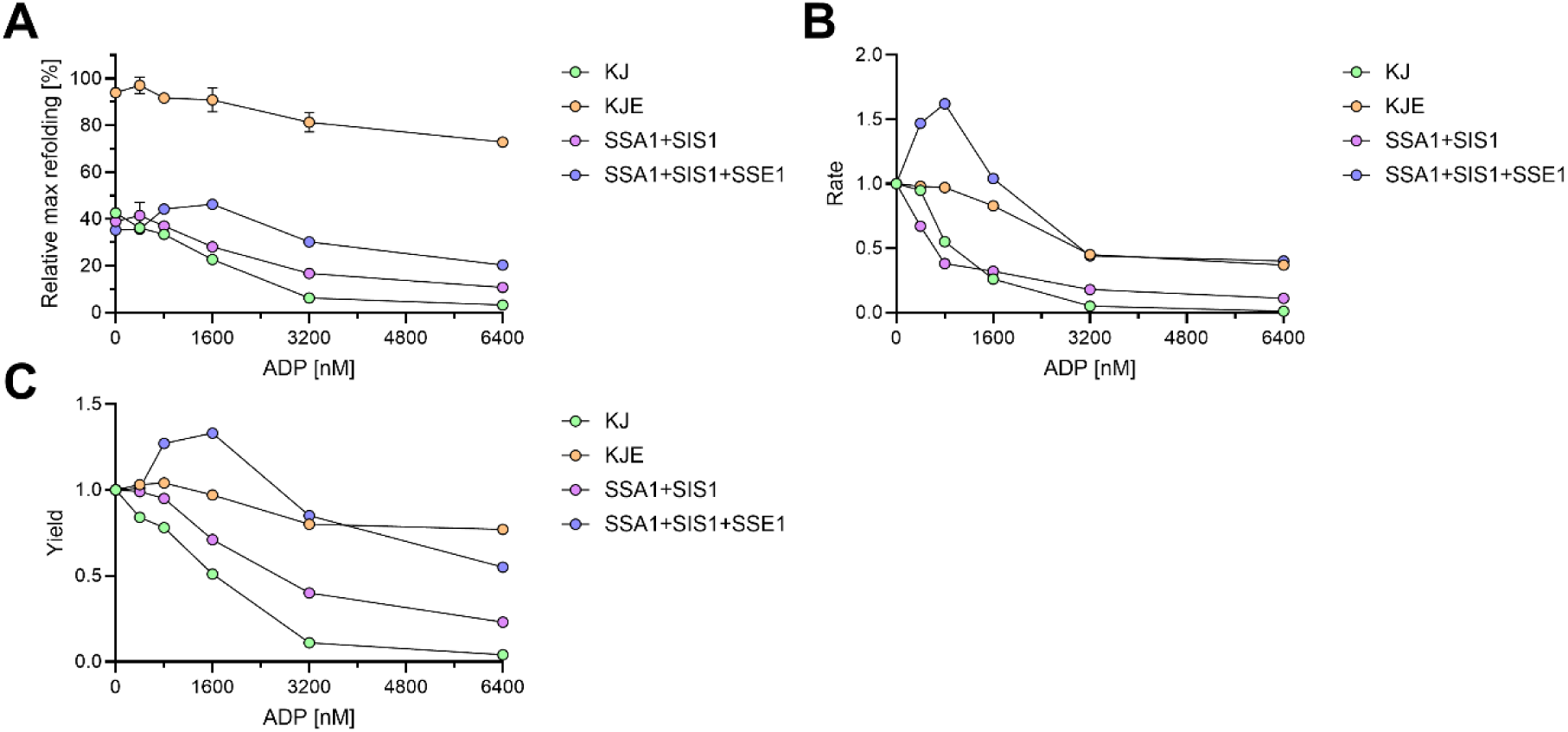
NEFs reduce inhibition by excess ADP of prokaryotic and eukaryotic Hsp70s. (**A**) 150 nM MlucV was pre-aggregated and incubated with 800 nM ATP, 6000 nM Ssa1 (or DnaK), 1000 nM Sis1 (or DnaJ) and increasing ADP, without or with 800 nM Sse1 (or GrpE). Recovered luciferase activity was measured after 3 hours. (**B**) Without and with Sse1, equimolar ADP respectively inhibits and activates Ssa1+JDP-mediated disaggregation and refolding of pre-aggregated MlucV. Optimal Luciferase refolding rates from pre-aggregated MlucV as in A, incubated at 30°C for up to 3 hours with 800 nM ATP, 6000 nM Ssa1, 1000 nM Sis1,1000 nM Ydj1 and increasing concentrations of ADP, without (blue), or with 800 nM Sse1. (**C**) Same as B but for the yields.

**Figure 7:**
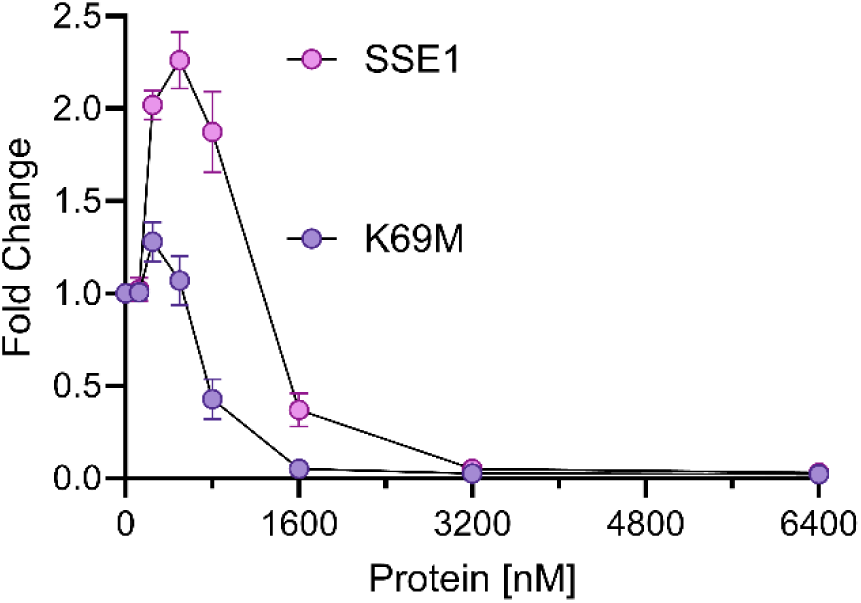
Fold change of refolding between Sse1 and Sse1-K69M. Fold change in refolding as a function of protein concentration for 6000 nM SSA and 1000 Sis1 that were added to pre-aggregated Luciferase (200 nM) with the addition of wild-type Sse1 or the K69M mutant after 2-hour.

### The role of ATP hydrolysis for Sse1 function

The Hsp110 co-chaperone family exhibits significant divergence from the Hsp70s from which it derived, particularly in the substrate-binding domain (SBD) and in the linker that in the Hsp70s specifically interacts with the HPD motive of J-domain co-chaperones. The SBDs and linkers of the Hsp110s also strongly repress ATP hydrolysis in their NBDs, more tightly than they do in Hsp70 (Vogel, Mayer et al. 2006, Swain, Dinler et al. 2007, Mayer 2018, Kumar, Peter et al. 2020). At variance with its Hsp70 ancestor, Sse1 has apparently lost the ability to interact with, and be activated by JDPs (as also predicted by AlphaFold protein complex predictions, Supplementary Figure 5), and compared to Ssa1, it has a greatly reduced ability to bind misfolded luciferase, both directly and indirectly, by way of JDP-mediated substrate-binding (Fig. 2A). In contrast, the perpetuation in evolution of its ATPase during two billion years since it stemmed out from the Hsp70s suggests that ATP hydrolysis still carries a central role in the ability of Sse1 to boost the basal stand-alone disaggregation mechanism of the Hsp70-JDP, even if it is not clear exactly under which condition it is necessary, as several studies showed contradictory results for the necessity of Hsp110 ATPase activity (Torrente and Shorter 2013). We compared the action of increasing concentrations of WT Sse1 to that of the known ATPase-defective Sse1-K69M mutant (Shaner, Trott et al. 2004, Raviol, Sadlish et al. 2006, Andréasson, Fiaux et al. 2008). Noticeably the isolated NBD of K69M, the ATPase deficient mutant, was found to hydrolyze ATP ∼15 times more slowly than the NBD of Sse1 (Kumar, Peter et al. 2020).

**Supplementary Figure 5:**
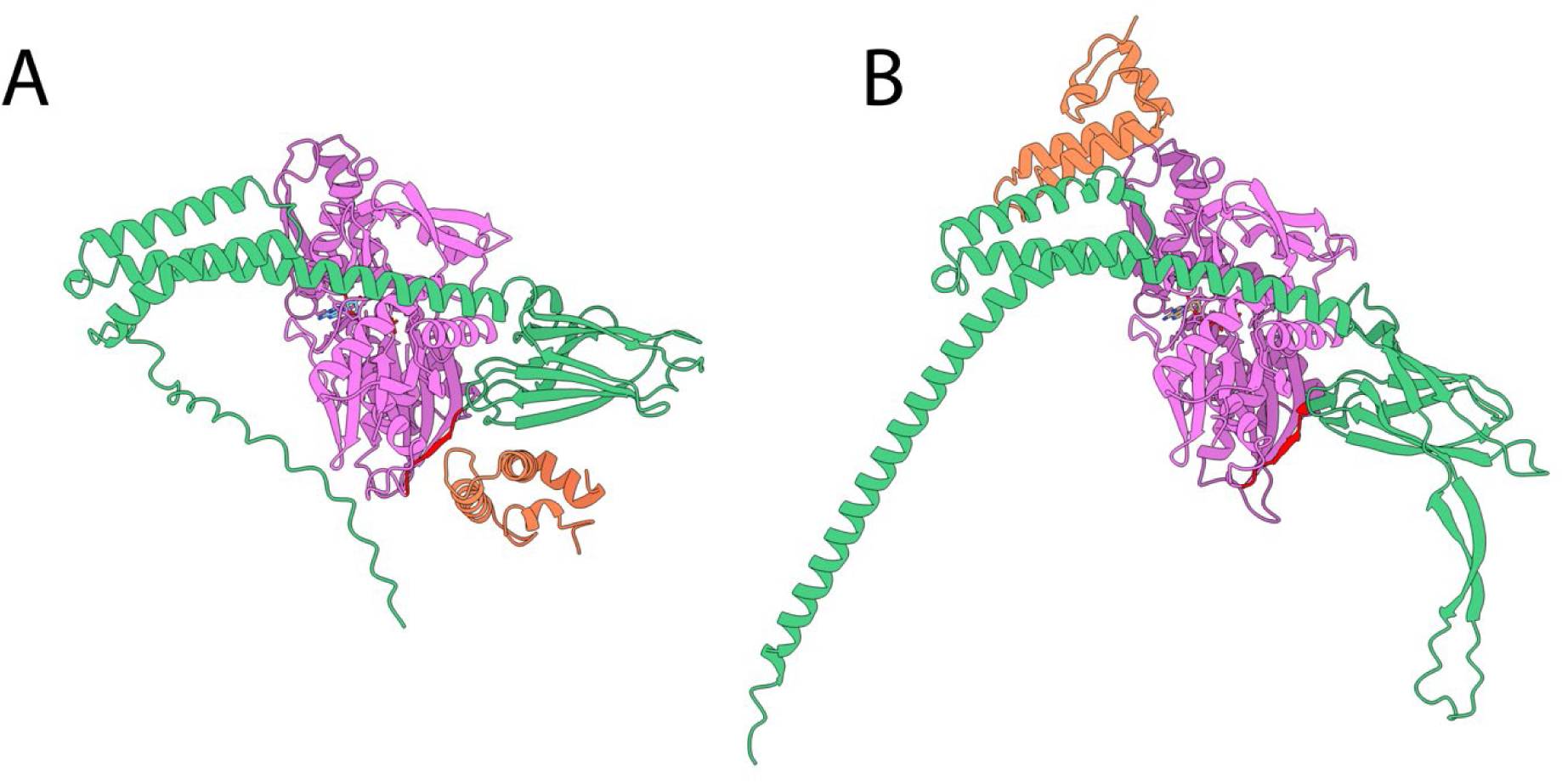
Sse1 has apparently lost the ability to interact with JDP. AlphaFold 3 best prediction of the ATP-Ssa1-Sis1 (**A**) and ATP-Sse1-Sis1 (**B**) complexes. NBDs in magenta, SBDs in green, linkers in red, J-domain of Sis1 in Orange.

Disaggregation assays showed that K69M activated significantly, even if only slightly, the Ssa1-driven disaggregation/refolding of pre-aggregated luciferase at an optimal concentration of 200 nM, which was 3 times lower than for WT Sse1 (600 nM) (Fig 7). Activation of disaggregation/refolding by Ssa1-Sis1 by 200 nM K69M was significant, yet still 5 times less effective as with twice more WT Sse1. Correspondingly, inhibition by excess K69M Sse1 occurred at a concentration four times lower than for WT Sse1, suggesting that K69M associates to Ssa1 more strongly, and dissociates from it more slowly, than WT sse1. (Fig. 7). These data hint that ATP hydrolysis being essential to drive an obligatory dissociation step of Sse1 from Ssa1, whose role is yet unclear for the effective Hsp70-JDP mediated disaggregation cycle.

### Disinhibition of SSE1’s ATPase does not correlate with the activation of disaggregation

The linker between the NBD and SBD domains of Hsp70 chaperones is highly conserved across the different kingdoms of life (DLLLLD in Ssa1, with relatively small variations, *e.g.* DVLLLD in DnaK)(Chakafana, Zininga et al. 2019). It specifically interacts with the highly conserved HPD motif of the J-domain in the crystal structure of the ATP-bound Hsp70-J-domain complex (Mayer and Gierasch 2019), while it should not interact with it in the ADP-bound Hsp70, when it is undocked from the NBD and completely extended and exposed to the interdomain space. The sequence of the linker is very different in Hsp110s (PFKFED in Sse1) (Chakafana, Mudau et al. 2021), and most likely does not interact with J-domains (Abramson, Adler et al. 2024), as also evidenced by the lack of Sse1’s ATPase activation by JDPs (Kumar, Peter et al. 2020) and by the lack of J-domain/NBD interaction in the AlphaFold 3 predicted structure of the Sse1/J-domain (of Sis1) complex (by comparison, the structure of the Ssa1/J-domain complex perfectly reproduces the DnaK/J-domain known complex) (Kityk, Kopp et al. 2018) (Sup. Fig. 5). The linker thus serves as a major hallmark distinguishing Hsp70s from Hsp110s (Swain, Dinler et al. 2007, Chakafana, Zininga et al. 2019). Furthermore, there is a complex interplay between the ATPase activity of Hsp70 and the domain interactions between NBD and SBD and with the linker: docking of the proximal part of the SBD’s α-helix on the NBD inhibits ATP hydrolysis, whereas docking of the flexible linker on the NBD accelerates ATP hydrolysis (English, Sherman et al. 2017).

We thus generated two Sse1 mutants that, according to the findings for DnaK, could change the ATPase activity of Sse1. In one mutant (Sse1-EAE), the PFKFED linker segment of Sse1 was changed into the DLLLLD linker from Ssa1 (Sup Fig. 2 and 3). We found that compared to WT Sse1, the ATPase activity of EAE (without substrate) was strongly enhanced (Fig. 8A), to a level comparable to the activity of Ssa1. Yet, at variance with Ssa1, EAE’s ATPase activity was equally high with and without Sis1. This indicates that the ATP hydrolysis is strongly repressed both by the PFKFED linker and by other parts of the SBD and becomes partially de-repressed when replaced by Hsp70’s linker segment. This also hints that the replacement of Hsp110’s linker by Hsp70’s did not suffice to restore Sse1’s ability to bind Sis1 and to further activate its ATPase, nor did it restore Sse1’s ability to bind and process aggregates. We then tested the ability of increasing doses of EAE to facilitate protein disaggregation by Ssa1 in the presence of Sis1 and found that it recapitulated the same behavior as WT Sse1, with sub-stoichiometric amounts optimally activating, and stoichiometric amount strongly inhibiting Ssa1-Sis1-mediated disaggregation and refolding (Fig. 8B). Thus, although more wasteful in ATP, EAE was as a potent co-disaggregase NEF as WT Sse1, and its mildly unleashed constitutive ATPase activity did not change the degree of inhibition by excess Sse1 binding to Ssa1.

**Figure 8:**
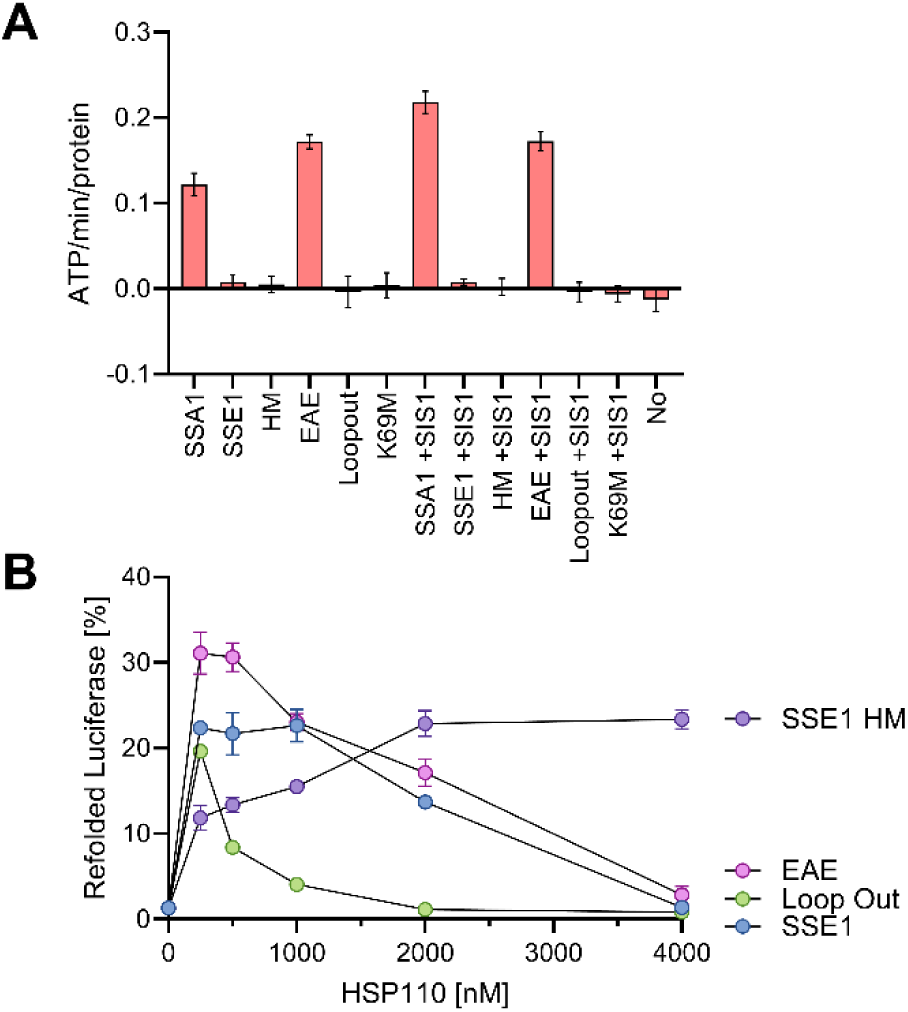
ATPase and activation profiles of Sse1 mutants on Ssa1-Sis1 disaggregation. **A**: The rate of ATP hydrolysis measured for 60 min at RT, in the absence or presence of 6000 nM Ssa1or WT Sse1, K69M, EAE, HM or Loopout mutants, in the absence or presence of 1000 nM Sis1. **B**. Luciferase refolding yields of 200 nM preaggregated Luciferase following incubation for 180 min at RT, in the presence of 6000 nM Ssa1, 1000 nM Sis1 and increasing concentrations of WT Sse1, K69M, EAE, HM or Loopout mutants.

To investigate the effect of the docking of other parts of Sse1’s SBD, of the α-helical lid in particular, on ATP hydrolysis in the NBD, we generated the mutant Sse1-HM in which the proximal part of the helical lid of Sse1 was exchanged for the corresponding helical segment from Ssa1 (Mayer 2018, Mayer and Gierasch 2019) (Sup Fig. 3). Since this segment did not co-evolve with Sse1’s NBD, we expected it to bind less tightly, and thereby possibly perturb the strong repression by the lid of the ATPase activity. But the results showed that ATP hydrolysis in the HM mutant remained as tightly auto repressed as in WT Sse1 (Fig. 8A), suggesting that other regions of the SBD contribute to Sse1’s ATPase autorepression. We tested the ability of increasing doses of HM mutant to facilitate protein disaggregation by Ssa1 in the presence of Sis1 and found that, unexpectedly, the ATPase remained as tightly auto-repressed as in WT Sse1 (Fig. 8A). In contrast, we found that the HM mutation apparently reduced the affinity of Sse1 for Ssa1 such that five times more HM mutant was necessary to half activate (EC_50_) the refolding reaction (500 nM instead of 100 nM), and equimolar HM mutant of did not inhibit the Ssa1-Sis1-mediated disaggregation and refolding reaction. This is suggesting that also the helical lid of Sse1 could be implicated in the regulation of its ATPase cycle, possibly coupling ATP hydrolysis in the NBD of Sse1 to a conformational change in its SBD (Kumar, Peter et al. 2020). This would be like the regulation of DnaK’s ATP hydrolysis by contacts between its NBD and helical lid (Vogel, Mayer et al. 2006, Swain, Dinler et al. 2007).

### The role of loop extensions in Sse1’s SBD

A hallmark of all Hsp110s is the presence of loop extensions with unclear, possibly unfolded structures, in the SBD β-basket and in the C-terminal end of the lid. Sse1 is no exception, although the length of the extra loops is very limited compared to plant and metazoan homologues. To explore the role of these extensions, we generated the Sse1 Loopout mutant, where the two loops have been removed (Sup Fig. 2 and 3). We tested the effect of the Loopout mutations on the intrinsic ATPase activity and the effect of increasing concentrations of Sse1 Loopout on its ability to boost disaggregation by Ssa1-Sis1. Because the Loopout mutant has a wild-type NBD, a canonical Sse1 linker and the extensions are not supposed to have interactions with the NBD, we expectedly found that the ATPase activity of Loopout was similarly very low, auto-repressed, as in the WT Sse1 (Fig. 8A). Surprisingly, the Loopout mutant was found to activate the disaggregation action of Ssa1-Sis1 similarly to the K69M hydrolysis-deficient mutant *i.e.* it enhanced disaggregation almost as for Sse1 albeit at smaller concentrations (up to 250 nM), and excess was inhibitory at lower concentrations than for WT Sse1, with an IC_50_ at about 500 nM (Fig. 8B). This is suggesting that to boost the Ssa1-JDP disaggregation cycle, Sse1’s extra-loops might favor, somehow, the obligatory dissociation from Ssa1of Sse1, that requires ATP hydrolysis by Sse1 and is otherwise tightly auto-repressed in the other phases of the cycle.

## Discussion

The presence of Hsp110 in the genome of all eukaryotes and their functional role raises two intersecting questions: i) what is their molecular mechanism of action? ii) What are the evolutionary steps that have transformed an ancestral Hsp70 into a Nucleotide Exchange Factor? We can break these questions into different sub-issues.

### Hsp70-Hsp110 interactions

Activation and inhibition of Hsp70 in a dose-dependent manner clearly hint at a direct interaction between Hsp70 and Hsp110. The contacts between the NBDs of the two proteins, as observed both in their crystal structures and in AI-generated predictions (AlphaFold 3) (Liu and Hendrickson 2007, Polier, Dragovic et al. 2008, Abramson, Adler et al. 2024) mirror the NBD-NBD arrangement found in the Hsp70 homodimer crystal structure (Sarbeng, Liu et al. 2015, Yang, Nune et al. 2015, Mayer 2021). Furthermore, the relevant residue-residue contacts in the Hsp70 homodimer have been also detected by co-evolutionary analysis (Malinverni, Marsili et al. 2015), and their perturbation causes defects in the function of DnaK (Sarbeng, Liu et al. 2015), strongly hinting at a functional role of the homodimer, conserved through evolution. We propose here that Hsp110s have evolved on this pre-existing Hsp70-Hsp70 interaction template. Given that GrpE and Sse1 interact at similar positions with the NBD of their corresponding Hsp70’s, and cause a comparable, strong decrease of ADP affinity, it is tempting to speculate that the transient formation of Hsp70 homodimers through their NBDs might accelerate ADP release and ATP rebinding. Such a concentration-dependent, self-nucleotide exchange activity could rationalize the significant, stand-alone unfolding, disaggregating and refolding activity that we observed, *in vitro*, with both prokaryotic and eukaryotic Hsp70s and their JDPs, in the absence of their dedicated NEFs. While the existence and functional importance of DnaK and Hsp70 homodimers have been demonstrated (Thompson, Bernard et al. 2012, Morgner, Schmidt et al. 2015, Sarbeng, Liu et al. 2015, Takakuwa, Nitika et al. 2019, Trcka, Durech et al. 2019), their precise molecular functional mechanism, and their possible evolution into functional Hsp70-Hsp110 heterodimers in eukaryotes, particularly effective for the disaggregation of large compact fibrils, remains to be further investigated.

### Hsp110 ATPase activity

The enhancement of Hsp70’s disaggregation and refolding activity occurs at low concentrations of Hsp110. This suggests that Hsp110 preferentially targets the limited pool of substrate-bound, ADP-bound Hsp70, rather than the excess of ATP-bound Hsp70 present in solution. Such specificity would imply that Hsp110 must have a very high affinity for the ADP-bound form of Hsp70—an affinity that must be rapidly downregulated after nucleotide exchange (which is certainly one, though likely not the only, function of Hsp110). This interpretation is supported by Alphafold 3 showing that ATP-Sse1 prefers ADP-bound, rather ATP-bound Ssa1, to form a heterodimer (Figure 1A) and by the data from the K69M mutant, which inhibits Hsp70 at lower concentrations, compared to WT Sse1. Indeed, a much slower ATPase would likely retain high-affinity even for ATP-bound Hsp70 in solution, and at high concentrations Sse1 (and even more so K69M) would form overly bulky complexes with Ssa1 in solution, whose size would hinder their binding to aggregates.

These observations suggest that the ATPase activity of Hsp110 plays a crucial role in modulating this affinity switch, to drive an obligatory step of dissociation of Sse1 from Ssa1, necessary for a new disaggregation cycle. From an evolutionary perspective, this proposal points to a functional repurposing—rather than a loss—of the ancestral Hsp70 ATPase, which has been conserved for over two billion years of Hsp110 evolution.

### Substrate (Hsp70) sensing

Just as JDP-locked ATP-Hsp70s “know” when to hydrolyze ATP through their in-built SBD sensor for substrate binding and through the interaction of their highly conserved linker with the HPD motif of J-domains, we would expect Hsp110 to only use its ATPase activity when useful, that is when Hsp70 undergoes nucleotide exchange. The EAE and Loopout mutants help shed light on this issue.

EAE had an intrinsic ATPase rate of 0.17 ATP min^−1^ that is much higher than the wild type Sse1, but their disaggregation/refolding activity was similar. It is tempting to propose that the wild-type linker has been specifically selected by evolution to avoid interaction with the HPD conserved triad of J-domains and eliminate a spurious interaction without affecting the rest of the function. The Loopout mutant, instead, had an effect on the action of Ssa1 similar to the ATPase-deficient K69M mutant (and similar to the mutant obtained by the deletion of the acidic extension of APG2 (Cabrera, Dublang et al. 2019), raising the possibility that in addition to loss of mass/volume, changes in the structural organization of Hsp70 could be sensed by the SBD of Hsp110 through the extensions, and allosterically be transmitted to the NBD to stimulate ATP hydrolysis and complex dissociation. It is remarkable that also in this case, evolution may not to have invented a new mechanism but rather have repurposed the pre-existing SBD-NBD sensing and allosteric communication.

### Role of Hsp110 as a NEF

While the NEF activity of Hsp110s has been already documented (Andréasson, Fiaux et al. 2008, Cyr 2008, Polier, Dragovic et al. 2008), here we have shown that it becomes even more relevant when ATP is limiting and the ADP concentrations increase, possibly become in excess over ATP. The presence of NEFs in general reduces the inhibition due to ADP and, in the case of Hsp110, can even favor the disaggregation reaction when, as in cells, hundreds of micromolar of ADP are present alongside millimolar of ATP. This NEF-driven increased resistance to the inhibition by high ADP concentrations is expected to be crucial in starving or quiescent bacteria and yeast, when ATP is not in excess, and NEFs may increase the affinity of Hsp70s for the increasingly rare ATP molecules.

### Molecular mechanism of action of Hsp110s

These considerations bring us back to the unresolved question of the true mechanism behind Hsp110’s action. If Hsp110 were simply nucleotide exchange factors (NEFs), as widely thought, following nucleotide exchange they would promote Hsp70 dissociation from the substrate, halting their disaggregation/unfolding process. Thus, their ability to enhance the action of Hsp70s seems counterintuitive. In previous work, the authors proposed a solution to this paradox, suggesting that the bulkiness of Hsp110s allows them to act as selective NEFs. Specifically, Hsp110s would target isolated, unproductive Hsp70s on aggregates, accessible due to the absence of steric hindrance from other nearby Hsp70s. By contrast, Hsp110s would not act on Hsp70 clusters, which were thought to be the disaggregation foci, due to the dense packing of Hsp70s in these regions. In this model, Hsp110s (and similarly bulky NEFs, such as BAG1 conjugated with large moieties) would free inactive Hsp70s, making them available for further cluster formation, or clear space around clusters, promoting their growth and thus their disaggregation efficiency. However, this model has a flaw: Hsp70 clusters necessarily form where steric hindrance is minimal, making these sites also favorable for Hsp110 binding. This raises a new perspective: we propose that, upon binding to substrate-bound Hsp70s, Hsp110s (or bulky BAG1) may increase entropic pulling forces, resulting in faster disaggregation. If the increase of the disaggregation rate is exponential (in the manner of Kramers’ theory (Rukes, Rebeaud et al. 2024)) and larger than the increase of the nucleotide exchange rate and consequent substrate dissociation, this might indeed lead to the observed enhancement by Hsp110 rather than by inhibition (Rohland, Kityk et al. 2022). Thus, in addition to being a NEF, merely accelerating ADP to ATP exchange, Hsp110 would boost the basal disaggregation activity of Hsp70s by increasing their effective size and enhancing entropic pulling strokes during the short time following ADP release from Hsp70’s NBDs, causing SBD opening and release of the substrate.

This effect increases with increasing Hsp110 concentration, as long as they are sufficiently small, because more of them bind to the small fraction of substrate-bound Hsp70s. At higher concentrations, however, Hsp110s would begin binding to free Hsp70s in solution, resulting in larger complexes with a reduced ability to bind to the aggregates. As a result, Hsp110 excessive binding would impede disaggregation at larger Hsp110 concentrations, leading to the observed inhibition.

Nucleotide Exchange Factors of the Hsp110 family represent an interesting evolutionary enigma. They belong to the Hsp70 family, but their presence in all eukaryotes with multiple cytosolic paralogs and one *endoplasmic reticulum* member suggests that their divergence and duplication preceded the appearance of the Last Eukaryotic Common Ancestor. Here we have highlighted how some of the transformations that have turned the Hsp70 disaggregating machinery into a NEF have likely resulted from the repurposing of pre-existent Hsp70 features, such as dimerization, ATPase activity and its triggering by interaction sensing and interdomain allosteric communication. We have in the end proposed a model for the molecular mechanism of action of Hsp110s as enhancers of entropic pulling forces.

More work is needed to decide whether this model, or the others that have been proposed (Wentink, Nillegoda et al. 2020), is the correct one, and, from an evolutionary perspective, more work is also needed to fully understand the role of the Hsp70 homodimer, which we have proposed here to be an ancestral template for the Hsp70-Hsp110 interaction.

## Methods

### Strains and plasmids

Wild-type *Ssa1*, *Sse1*, *Sis1*, *Ydj1, and* mutants were cloned in the pE-SUMO vector for propagation in *E. coli.* BL21-CodonPlus (DE3)-RIPL.

### Generating mutant proteins

Point mutations were introduced through site-directed mutagenesis PCR using Q5 Site-Directed Mutagenesis Kit (New England Biolabs, cat. no. E0554). Mutants were constructed by PCR by amplifying the selected regions with the primers list from Supplementary Table 2, and proteins used for this study are listed in Supplementary Table 3.

### Proteins purification

For purification of the His10-SUMO tagged wild-type Sse1, Ssa1, Sis1, Ydj1, and mutants, were expressed and purified from *E. coli* BL21-CodonPlus (DE3)-RIPL cells with IPTG induction (final 0.5 mM for Ssa1 and Sse1 and 0.2 for Ydj1, Sis1, and mutants) at 18 °C, overnight. Briefly, cells were grown in LB medium + ampicillin at 37 °C to OD600 ∼0.4-0.5. Protein expression was induced by the addition of 0.5 mM IPTG for 3 hours. Cells were harvested, washed with chilled PBS, and resuspended in buffer A (20 mM Tris-HCl pH 7.5, 150 mM KCl, 5% glycerol, 2mM DTT, 20mM MgCl2) containing 5 mM imidazole, 1mg/ml Lysozyme, 1mM PMSF for 1 h. Cells were lysed by sonication. After high-speed centrifugation (16000 rpm, 30 min/4°C), the supernatant was loaded onto a gravity flow-based Ni-NTA metal affinity column (2 ml beads, cOmplete His-Tag Purification Resin from Merck), equilibrated and washed with 10 column volumes of buffer A containing 5 mM imidazole. After several washes with high salt buffer A (+150 mM KCl, 20mM Imidazole and 5 mM ATP), N-terminal His10-SUMO (small ubiquitin-related modifier) Smt3 tag was cleaved with Ulp1 protease (2mg/ml, 300 μl, added to beads with buffer (20 mM Tris-HCl pH 7.5, 150mM KCl, 10mM MgCl2, 5% glycerol, 2mM DTT). Digestion of His10 Smt3 was performed on the Ni-NTA resin by His6-Ulp1 protease. Because of dual His tags, His6-Ulp1 and His10-SUMO display a high affinity for Ni-NTA resin and remain bound to it during cleavage reaction. After overnight digestion at 4°C, the unbound fraction is collected (which contains only the native proteins). Proteins were further purified by concentrating to ∼3mg/ml and applying to a size exclusion column (Superdex-200 increase, 10/30 GE Healthcare) equilibrated in buffer A containing 5 mM ATP. Pure fractions were pooled, concentrated by ultrafiltration using Amicon Ultra (Millipore), aliquoted, and stored at −80 °C. All protein concentrations were determined spectrophotometrically at 562 nm using BCA Protein Assay Kit− Reducing Agent Compatible (cat no. 23250). The purified proteins were collected, concentrated, and stored at −80°C for further use.

### Choice of substrates: Luciferase and MlucV

Two well-established chaperone substrates were used: stably preformed luciferase aggregates (Shorter-like aggregate (Shorter 2011)) and stably urea-preformed MlucV aggregates (Tiwari, Fauvet et al. 2023). MlucV is composed of a firefly luciferase flanked by two domains of GFP-derived fluorophores, whose fluorescent resonance energy transfer (FRET) values inform on the different states of the reported protein: already stably aggregated, transiently unfolded or native. It allows following the conversion by the Hsp70-JDP-Hsp110 chaperone system of stably pre-aggregated substrates formed prior chaperone exposure, to completely or partially unfolded intermediates and finally to the natively refolded products of the chaperone action (Tiwari, Fauvet et al. 2023). Recovered luciferase activity was used to confirm that the chaperone did mediate polypeptide disaggregation and unfolding that subsequently lead to the native refolding of the core luciferase domain in the preaggregated MlucV.

### FRET Measurements and Proximity Ratio Calculation

Ensemble relative FRET ratios were determined by measuring the maximum fluorescence emission intensities of the donor (ED) and acceptor (EA) fluorophores upon excitation of the donor at 405 nm, as described previously (Fritz, Letzelter et al. 2013, Wood, Ormsby et al. 2018). The fluorescence emission spectra of the MlucV reporter were recorded using a Tecan SPARK (Männedorf, Switzerland) in a 96-well microtiter plate. For donor fluorophore, excitation and emission wavelengths were at 405nm and 493nm with an excitation bandwidth of 20 nm. For acceptor, excitation and emission wavelengths were at 405nm and 525nm with an excitation bandwidth of 20 nm. To minimize direct excitation of the acceptor, donor excitation was strictly maintained at 405 nm. Background subtraction was performed using spectra obtained from buffer-only samples. Where applicable, baseline-corrected spectra were further normalized to their respective areas. All measurements were carried out in LRB refolding buffer (20 mM HEPES-KOH, pH 7.5, 150 mM KCl, 10 mM MgCl₂) supplemented with 4 mM ATP, unless stated otherwise. Each experiment was performed in triplicate or more to ensure reproducibility. Normalized FRET ratios relative to that of native MlucV were calculated as follows:

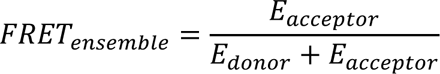

Normalized FRET efficiencies relative to that of native MlucV were calculated as follows (Kopp, Nowak et al. 2018, Tiwari, Fauvet et al. 2023):

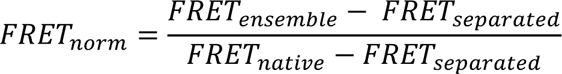

where FRET_ensemble_ is the measured ensemble FRET efficiency, FRET_separated_ is the calculated ensemble FRET measured in a solution of separated mTFP1 and Venus (∼0.335) and FRET_native_ is the measured ensemble FRET of native MlucV (∼0.425).

A FRET value for the native MlucV set to be 100%, a FRET value for the unconnected fluorophores set to be 0%. FRET values for the stable, pre-aggregated MlucV (130-170% depending on aggregation conditions) and finally, a FRET value for the unfolded luciferase in MlucV the presence of 4.8M urea (found to be 40% of native), and for the chaperone-bound unfolded luciferase in MlucV (found to be generally less than native).

Thus, FRET measures allowed to follow the ATP-dependent conversion of stably aggregated polypeptide substrates formed prior chaperone exposure, to completely or partially unfolded chaperone-bound intermediates and finally, to the natively refolded products of the chaperone reaction (Tiwari, Fauvet et al. 2023). Recovered luciferase activity was used to confirm that the chaperone did mediate polypeptide disaggregation and unfolding that subsequently lead to the native refolding of the core luciferase domain in the preaggregated MlucV. For comparison with the MlucV results, urea-pre aggregated firefly luciferase was also used as a chaperone substrate as (Shorter 2011) and only the luciferase activity, without FRET was used to assess the chaperone disaggregation and refolding activities.

### Luciferase and MlucV refolding assay

Luciferase and MlucV activity were measured as described previously (Bischofberger, Han et al. 2003, Sharma, De los Rios et al. 2010, Tiwari, Fauvet et al. 2023). In the presence of oxygen, luciferase catalyzes the conversion of D-luciferin and ATP into oxyluciferin, CO2, AMP, PPi, and hν. Generated photons were counted with a Victor Light 1420 Luminescence Counter from Perkin-Elmer (Turku, Finland) in a 96-well microtiter plate format.

### ATPase assay (Malachite Green)

Colorimetric determination of Pi produced by ATP hydrolysis was performed using the Malachite Green Assay Kit (Sigma-Aldrich, Switzerland) and as described previously (Lee, Roh et al. 2019). Several concentrations of Hsp70 (Ssa1), JDPs (Ydj1, Sis1) or mutants were mixed with or without substrate (200 nM of pre-aggregated luciferase) and with 1 mM of ATP and incubated for 1 hour at 25°C. 8 µL of each sample was taken and put inside a 96-Well plate with 72 µL of H2O. A 20-µl volume of Malachite Green reaction buffer was added, and the samples were mixed thoroughly and incubated at 25°C for 30 min before measuring at 620nm on a plate reader (Tecan SPARK, Männedorf, Switzerland). The rate of intrinsic ATP hydrolysis was deduced by subtracting the signal from ATP in the absence of a chaperone.

### Phylogenetic tree

Maximum likelihood phylogenetic trees were generated by MEGA X (Kumar, Stecher et al. 2018), using JTT distance matrix and NJ/BioNJ initial tree. Trees were rooted at midpoint and made with iTOL (Letunic and Bork 2021).

## Author contributions

Conceptualization: MER, PG, PDLR; Methodology: PG, PDLR, MER, BF; Investigation: PG, PDLR, MER; Visualization: PG, PDLR, MER, BF; Supervision: PG, PDLR; Writing—original draft: PG, PDLR, MER; Writing—review & editing: PG, PDLR, MER.

## Supplementary Tables

**Supplementary Table 2.**
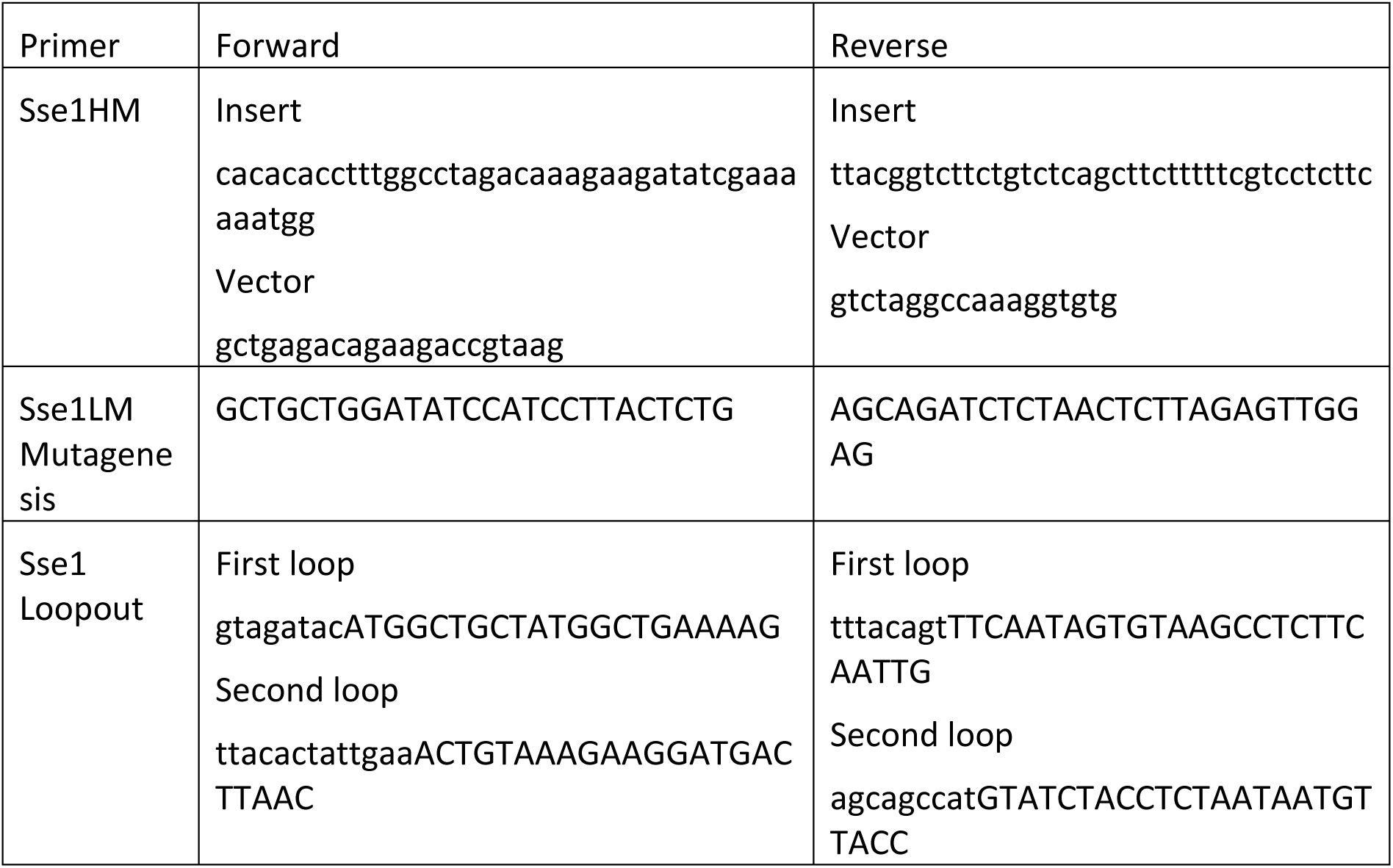
Primers used in this study.

**Supplementary Table 3.** Proteins used in this study.

| Protein | Protein family | References |
| --- | --- | --- |
| Ssa1 | Hsp70 WT | (Werner-Washburne, Stone et al. 1987) |
| Sse1 | Hsp110 WT | (Mukai, Kuno et al. 1993) |
| Sse2 | Hsp110 WT | (Mukai, Kuno et al. 1993) |
| Sis1 | JDP WT | (Luke, Sutton et al. 1991) |
| Ydj1 | JDP WT | (Caplan and Douglas 1991) |
| YS | JDP Swap | (Rebeaud, Tiwari et al. 2024) |
| Sse1HM | Hsp110 mutant | This study |
| Sse1LM (EAE) | Hsp110 mutant | This study |
| Sse1 Loopout | Hsp110 mutant | This study |
| Sse1 K69M | Hsp110 mutant | (Raviol, Sadlish et al. 2006) |

## References

Abramson, J., J. Adler, J. Dunger, R. Evans, T. Green, A. Pritzel, O. Ronneberger, L. Willmore, A. J. Ballard, J. Bambrick, S. W. Bodenstein, D. A. Evans, C.-C. Hung, M. O’Neill, D. Reiman, K. Tunyasuvunakool, Z. Wu, A. Žemgulytė, E. Arvaniti, C. Beattie, O. Bertolli, A. Bridgland, A. Cherepanov, M. Congreve, A. I. Cowen-Rivers, A. Cowie, M. Figurnov, F. B. Fuchs, H. Gladman, R. Jain, Y. A. Khan, C. M. R. Low, K. Perlin, A. Potapenko, P. Savy, S. Singh, A. Stecula, A. Thillaisundaram, C. Tong, S. Yakneen, E. D. Zhong, M. Zielinski, A. Žídek, V. Bapst, P. Kohli, M. Jaderberg, D. Hassabis and J. M. Jumper (2024). “Accurate structure prediction of biomolecular interactions with AlphaFold 3.” Nature 630(8016): 493–500.

Andréasson, C., J. Fiaux, H. Rampelt, S. Druffel-Augustin and B. Bukau (2008). “Insights into the structural dynamics of the Hsp110–Hsp70 interaction reveal the mechanism for nucleotide exchange activity.” Proceedings of the National Academy of Sciences 105(43): 16519–16524.

Bischofberger, P., W. Han, B. Feifel, H.-J. Schönfeld and P. Christen (2003). “d-Peptides as Inhibitors of the DnaK/DnaJ/GrpE Chaperone System*.” Journal of Biological Chemistry 278(21): 19044–19047.

Bork, P., C. Sander and A. Valencia (1992). “An ATPase domain common to prokaryotic cell cycle proteins, sugar kinases, actin, and hsp70 heat shock proteins.” Proc Natl Acad Sci U S A 89(16): 7290–7294.

Bracher, A. and J. Verghese (2015). “The nucleotide exchange factors of Hsp70 molecular chaperones.” Front Mol Biosci 2: 10.

Brownridge, P., C. Lawless, A. B. Payapilly, K. Lanthaler, S. W. Holman, V. M. Harman, C. M. Grant, R. J. Beynon and S. J. Hubbard (2013). “Quantitative analysis of chaperone network throughput in budding yeast.” Proteomics 13(8): 1276–1291.

Cabrera, Y., L. Dublang, J. A. Fernández-Higuero, D. Albesa-Jové, M. Lucas, A. R. Viguera, M. E. Guerin, J. M. G. Vilar, A. Muga and F. Moro (2019). “Regulation of Human Hsc70 ATPase and Chaperone Activities by Apg2: Role of the Acidic Subdomain.” Journal of Molecular Biology 431(2): 444–461.

Caplan, A. J. and M. G. Douglas (1991). “Characterization of YDJ1: a yeast homologue of the bacterial dnaJ protein.” J Cell Biol 114(4): 609–621.

Chakafana, G., P. T. Mudau, T. Zininga and A. Shonhai (2021). “Characterisation of a unique linker segment of the Plasmodium falciparum cytosol localised Hsp110 chaperone.” International Journal of Biological Macromolecules 180: 272–285.

Chakafana, G., T. Zininga and A. Shonhai (2019). “The Link That Binds: The Linker of Hsp70 as a Helm of the Protein’s Function.” Biomolecules 9(10): 543.

Cyr, D. M. (2008). “Swapping Nucleotides, Tuning Hsp70.” Cell 133(6): 945–947.

De Los Rios, P. and A. Barducci (2014). “Hsp70 chaperones are non-equilibrium machines that achieve ultra-affinity by energy consumption.” Elife 3: e02218.

De Los Rios, P., A. Ben-Zvi, O. Slutsky, A. Azem and P. Goloubinoff (2006). “Hsp70 chaperones accelerate protein translocation and the unfolding of stable protein aggregates by entropic pulling.” Proceedings of the National Academy of Sciences 103(16): 6166–6171.

Doello, S., M. Burkhardt and K. Forchhammer (2021). “The essential role of sodium bioenergetics and ATP homeostasis in the developmental transitions of a cyanobacterium.” Current Biology 31(8): 1606–1615.e1602.

English, C. A., W. Sherman, W. Meng and L. M. Gierasch (2017). “The Hsp70 interdomain linker is a dynamic switch that enables allosteric communication between two structured domains.” J Biol Chem 292(36): 14765–14774.

Fauvet, B., A. Finka, M. P. Castanie-Cornet, A. M. Cirinesi, P. Genevaux, M. Quadroni and P. Goloubinoff (2021). “Bacterial Hsp90 Facilitates the Degradation of Aggregation-Prone Hsp70-Hsp40 Substrates.” Front Mol Biosci 8: 653073.

Fritz, R. D., M. Letzelter, A. Reimann, K. Martin, L. Fusco, L. Ritsma, B. Ponsioen, E. Fluri, S. Schulte-Merker, J. van Rheenen and O. Pertz (2013). “A versatile toolkit to produce sensitive FRET biosensors to visualize signaling in time and space.” Sci Signal 6(285): rs12.

Genevaux, P., C. Georgopoulos and W. L. Kelley (2007). “The Hsp70 chaperone machines of Escherichia coli: a paradigm for the repartition of chaperone functions.” Molecular Microbiology 66(4): 840–857.

Ghaemmaghami, S., W. K. Huh, K. Bower, R. W. Howson, A. Belle, N. Dephoure, E. K. O’Shea and J. S. Weissman (2003). “Global analysis of protein expression in yeast.” Nature 425(6959): 737–741.

Goeckeler, J. L., A. P. Petruso, J. Aguirre, C. C. Clement, G. Chiosis and J. L. Brodsky (2008). “The yeast Hsp110, Sse1p, exhibits high-affinity peptide binding.” FEBS Lett 582(16): 2393–2396.

Goloubinoff, P., A. S. Sassi, B. Fauvet, A. Barducci and P. De Los Rios (2018). “Chaperones convert the energy from ATP into the nonequilibrium stabilization of native proteins.” Nat Chem Biol 14(4): 388–395.

Imamoglu, R., D. Balchin, M. Hayer-Hartl and F. U. Hartl (2020). “Bacterial Hsp70 resolves misfolded states and accelerates productive folding of a multi-domain protein.” Nature Communications 11(1): 365.

Jumper, J., R. Evans, A. Pritzel, T. Green, M. Figurnov, O. Ronneberger, K. Tunyasuvunakool, R. Bates, A. Zidek, A. Potapenko, A. Bridgland, C. Meyer, S. A. A. Kohl, A. J. Ballard, A. Cowie, B. Romera-Paredes, S. Nikolov, R. Jain, J. Adler, T. Back, S. Petersen, D. Reiman, E. Clancy, M. Zielinski, M. Steinegger, M. Pacholska, T. Berghammer, S. Bodenstein, D. Silver, O. Vinyals, A. W. Senior, K. Kavukcuoglu, P. Kohli and D. Hassabis (2021). “Highly accurate protein structure prediction with AlphaFold.” Nature 596(7873): 583–589.

Kampinga, H. H., C. Andreasson, A. Barducci, M. E. Cheetham, D. Cyr, C. Emanuelsson, P. Genevaux, J. E. Gestwicki, P. Goloubinoff, J. Huerta-Cepas, J. Kirstein, K. Liberek, M. P. Mayer, K. Nagata, N. B. Nillegoda, P. Pulido, C. Ramos, P. De Los Rios, S. Rospert, R. Rosenzweig, C. Sahi, M. Taipale, B. Tomiczek, R. Ushioda, J. C. Young, R. Zimmermann, A. Zylicz, M. Zylicz, E. A. Craig and J. Marszalek (2019). “Function, evolution, and structure of J-domain proteins.” Cell Stress Chaperones 24(1): 7–15.

Kityk, R., J. Kopp and M. P. Mayer (2018). “Molecular Mechanism of J-Domain-Triggered ATP Hydrolysis by Hsp70 Chaperones.” Mol Cell 69(2): 227–237 e224.

Kopp, M. C., P. R. Nowak, N. Larburu, C. J. Adams and M. M. U. Ali (2018). “In vitro FRET analysis of IRE1 and BiP association and dissociation upon endoplasmic reticulum stress.” eLife 7: e30257.

Kumar, S., G. Stecher, M. Li, C. Knyaz and K. Tamura (2018). “MEGA X: Molecular Evolutionary Genetics Analysis across Computing Platforms.” Molecular Biology and Evolution 35(6): 1547–1549.

Kumar, V., J. J. Peter, A. Sagar, A. Ray, M. P. Jha, M. E. Rebeaud, S. Tiwari, P. Goloubinoff, F. Ashish and K. Mapa (2020). “Interdomain communication suppressing high intrinsic ATPase activity of Sse1 is essential for its co-disaggregase activity with Ssa1.” The FEBS Journal 287(4): 671–694.

Lawless, C., S. W. Holman, P. Brownridge, K. Lanthaler, V. M. Harman, R. Watkins, D. E. Hammond, R. L. Miller, P. F. Sims, C. M. Grant, C. E. Eyers, R. J. Beynon and S. J. Hubbard (2016). “Direct and Absolute Quantification of over 1800 Yeast Proteins via Selected Reaction Monitoring.” Mol Cell Proteomics 15(4): 1309–1322.

Lee, S., S. H. Roh, J. Lee, N. Sung, J. Liu and F. T. F. Tsai (2019). “Cryo-EM Structures of the Hsp104 Protein Disaggregase Captured in the ATP Conformation.” Cell reports 26(1): 29–36.e23.

Letunic, I. and P. Bork (2021). “Interactive Tree Of Life (iTOL) v5: an online tool for phylogenetic tree display and annotation.” Nucleic Acids Research 49(W1): W293–W296.

Liu, Q. and W. A. Hendrickson (2007). “Insights into Hsp70 chaperone activity from a crystal structure of the yeast Hsp110 Sse1.” Cell 131(1): 106–120.

Lu, Z. and D. M. Cyr (1998). “Protein folding activity of Hsp70 is modified differentially by the hsp40 co-chaperones Sis1 and Ydj1.” J Biol Chem 273(43): 27824–27830.

Luke, M. M., A. Sutton and K. T. Arndt (1991). “Characterization of SIS1, a Saccharomyces cerevisiae homologue of bacterial dnaJ proteins.” J Cell Biol 114(4): 623–638.

Mackenzie, R. J., C. Lawless, S. W. Holman, K. Lanthaler, R. J. Beynon, C. M. Grant, S. J. Hubbard and C. E. Eyers (2016). “Absolute protein quantification of the yeast chaperome under conditions of heat shock.” Proteomics 16(15-16): 2128–2140.

Malinverni, D., S. Marsili, A. Barducci and P. De Los Rios (2015). “Large-Scale Conformational Transitions and Dimerization Are Encoded in the Amino-Acid Sequences of Hsp70 Chaperones.” PLOS Computational Biology 11(6): e1004262.

Manalastas-Cantos, K., K. R. Adoni, M. Pfeifer, B. Martens, K. Grunewald, K. Thalassinos and M. Topf (2024). “Modeling Flexible Protein Structure With AlphaFold2 and Crosslinking Mass Spectrometry.” Mol Cell Proteomics 23(3): 100724.

Mayer, M. P. (2018). “Intra-molecular pathways of allosteric control in Hsp70s.” Philosophical Transactions of the Royal Society B: Biological Sciences 373(1749): 20170183.

Mayer, M. P. (2021). “The Hsp70-Chaperone Machines in Bacteria.” Frontiers in Molecular Biosciences 8.

Mayer, M. P. and B. Bukau (2005). “Hsp70 chaperones: cellular functions and molecular mechanism.” Cell Mol Life Sci 62(6): 670–684.

Mayer, M. P. and L. M. Gierasch (2019). “Recent advances in the structural and mechanistic aspects of Hsp70 molecular chaperones.” Journal of Biological Chemistry 294(6): 2085–2097.

Morgner, N., C. Schmidt, V. Beilsten-Edmands, I.-o. Ebong, Nisha A. Patel, Eugenia M. Clerico, E. Kirschke, S. Daturpalli, Sophie E. Jackson, D. Agard and Carol V. Robinson (2015). “Hsp70 Forms Antiparallel Dimers Stabilized by Post-translational Modifications to Position Clients for Transfer to Hsp90.” Cell Reports 11(5): 759–769.

Mukai, H., T. Kuno, H. Tanaka, D. Hirata, T. Miyakawa and C. Tanaka (1993). “Isolation and characterization of SSE1 and SSE2, new members of the yeast HSP70 multigene family.” Gene 132(1): 57–66.

Polier, S., Z. Dragovic, F. U. Hartl and A. Bracher (2008). “Structural Basis for the Cooperation of Hsp70 and Hsp110 Chaperones in Protein Folding.” Cell 133(6): 1068–1079.

Raviol, H., B. Bukau and M. P. Mayer (2006). “Human and yeast Hsp110 chaperones exhibit functional differences.” FEBS Lett 580(1): 168–174.

Raviol, H., H. Sadlish, F. Rodriguez, M. P. Mayer and B. Bukau (2006). “Chaperone network in the yeast cytosol: Hsp110 is revealed as an Hsp70 nucleotide exchange factor.” EMBO J 25(11): 2510–2518.

Rebeaud, M. E., S. Mallik, P. Goloubinoff and D. S. Tawfik (2021). “On the evolution of chaperones and cochaperones and the expansion of proteomes across the Tree of Life.” Proc Natl Acad Sci U S A 118(21).

Rebeaud, M. E., S. Tiwari, B. Fauvet, A. Mohr, P. Goloubinoff and P. De Los Rios (2024). “Autorepression of yeast Hsp70 cochaperones by intramolecular interactions involving their J-domains.” Cell Stress Chaperones 29(2): 338–348.

Richards, T. A., L. Eme, J. M. Archibald, G. Leonard, S. M. Coelho, A. de Mendoza, C. Dessimoz, P. Dolezal, L. K. Fritz-Laylin, T. Gabaldón, V. Hampl, G. J. P. L. Kops, M. M. Leger, P. Lopez-Garcia, J. O. McInerney, D. Moreira, S. A. Muñoz-Gómez, D. J. Richter, I. Ruiz-Trillo, A. E. Santoro, A. Sebé-Pedrós, B. Snel, C. W. Stairs, E. C. Tromer, J. J. E. van Hooff, B. Wickstead, T. A. Williams, A. J. Roger, J. B. Dacks and J. G. Wideman (2024). “Reconstructing the last common ancestor of all eukaryotes.” PLOS Biology 22(11): e3002917.

Rohland, L., R. Kityk, L. Smalinskaitė and M. P. Mayer (2022). “Conformational dynamics of the Hsp70 chaperone throughout key steps of its ATPase cycle.” Proceedings of the National Academy of Sciences 119(48): e2123238119.

Rosenzweig, R., N. B. Nillegoda, M. P. Mayer and B. Bukau (2019). “The Hsp70 chaperone network.” Nat Rev Mol Cell Biol 20(11): 665–680.

Rukes, V., M. E. Rebeaud, L. W. Perrin, P. De Los Rios and C. Cao (2024). “Single-molecule evidence of Entropic Pulling by Hsp70 chaperones.” Nat Commun 15(1): 8604.

Russell, R., A. Wali Karzai, A. F. Mehl and R. McMacken (1999). “DnaJ dramatically stimulates ATP hydrolysis by DnaK: insight into targeting of Hsp70 proteins to polypeptide substrates.” Biochemistry 38(13): 4165–4176.

Sarbeng, E. B., Q. Liu, X. Tian, J. Yang, H. Li, J. L. Wong, L. Zhou and Q. Liu (2015). “A Functional DnaK Dimer Is Essential for the Efficient Interaction with Hsp40 Heat Shock Protein *.” Journal of Biological Chemistry 290(14): 8849–8862.

Shaner, L., R. Sousa and K. A. Morano (2006). “Characterization of Hsp70 Binding and Nucleotide Exchange by the Yeast Hsp110 Chaperone Sse1.” Biochemistry 45(50): 15075–15084.

Shaner, L., A. Trott, J. L. Goeckeler, J. L. Brodsky and K. A. Morano (2004). “The function of the yeast molecular chaperone Sse1 is mechanistically distinct from the closely related hsp70 family.” J Biol Chem 279(21): 21992–22001.

Sharma, S. K., P. De los Rios, P. Christen, A. Lustig and P. Goloubinoff (2010). “The kinetic parameters and energy cost of the Hsp70 chaperone as a polypeptide unfoldase.” Nat Chem Biol 6(12): 914–920.

Sharma, S. K., P. De Los Rios and P. Goloubinoff (2011). “Probing the different chaperone activities of the bacterial HSP70-HSP40 system using a thermolabile luciferase substrate.” Proteins: Structure, Function, and Bioinformatics 79(6): 1991–1998.

Shorter, J. (2011). “The mammalian disaggregase machinery: Hsp110 synergizes with Hsp70 and Hsp40 to catalyze protein disaggregation and reactivation in a cell-free system.” PLoS One 6(10): e26319.

Sontag, E. M., R. S. Samant and J. Frydman (2017). “Mechanisms and Functions of Spatial Protein Quality Control.” Annu Rev Biochem 86: 97–122.

Swain, J. F., G. Dinler, R. Sivendran, D. L. Montgomery, M. Stotz and L. M. Gierasch (2007). “Hsp70 Chaperone Ligands Control Domain Association via an Allosteric Mechanism Mediated by the Interdomain Linker.” Molecular Cell 26(1): 27–39.

Takaine, M., H. Imamura and S. Yoshida (2022). “High and stable ATP levels prevent aberrant intracellular protein aggregation in yeast.” eLife 11: e67659.

Takakuwa, J. E., Nitika, L. E. Knighton and A. W. Truman (2019). “Oligomerization of Hsp70: Current Perspectives on Regulation and Function.” Front Mol Biosci 6: 81.

Thompson, A. D., S. M. Bernard, G. Skiniotis and J. E. Gestwicki (2012). “Visualization and functional analysis of the oligomeric states of Escherichia coli heat shock protein 70 (Hsp70/DnaK).” Cell Stress Chaperones 17(3): 313–327.

Tiwari, S., B. Fauvet, S. Assenza, P. De Los Rios and P. Goloubinoff (2023). “A fluorescent multi-domain protein reveals the unfolding mechanism of Hsp70.” Nat Chem Biol 19(2): 198–205.

Torrente, M. P. and J. Shorter (2013). “The metazoan protein disaggregase and amyloid depolymerase system: Hsp110, Hsp70, Hsp40, and small heat shock proteins.” Prion 7(6): 457–463.

Trcka, F., M. Durech, P. Vankova, J. Chmelik, V. Martinkova, J. Hausner, A. Kadek, J. Marcoux, T. Klumpler, B. Vojtesek, P. Muller and P. Man (2019). “Human Stress-inducible Hsp70 Has a High Propensity to Form ATP-dependent Antiparallel Dimers That Are Differentially Regulated by Cochaperone Binding*.” Molecular & Cellular Proteomics 18(2): 320–337.

Vogel, M., M. P. Mayer and B. Bukau (2006). “Allosteric Regulation of Hsp70 Chaperones Involves a Conserved Interdomain Linker*.” Journal of Biological Chemistry 281(50): 38705–38711.

Wentink, A. S., N. B. Nillegoda, J. Feufel, G. Ubartaitė, C. P. Schneider, P. De Los Rios, J. Hennig, A. Barducci and B. Bukau (2020). “Molecular dissection of amyloid disaggregation by human HSP70.” Nature 587(7834): 483–488.

Werner-Washburne, M., D. E. Stone and E. A. Craig (1987). “Complex interactions among members of an essential subfamily of hsp70 genes in Saccharomyces cerevisiae.” Mol Cell Biol 7(7): 2568–2577.

Wood, R. J., A. R. Ormsby, M. Radwan, D. Cox, A. Sharma, T. Vopel, S. Ebbinghaus, M. Oliveberg, G. E. Reid, A. Dickson and D. M. Hatters (2018). “A biosensor-based framework to measure latent proteostasis capacity.” Nat Commun 9(1): 287.

Wyszkowski, H., A. Janta, W. Sztangierska, I. Obuchowski, T. Chamera, A. Klosowska and K. Liberek (2021). “Class-specific interactions between Sis1 J-domain protein and Hsp70 chaperone potentiate disaggregation of misfolded proteins.” Proc Natl Acad Sci U S A 118(49): e2108163118.

Yang, J., M. Nune, Y. Zong, L. Zhou and Q. Liu (2015). “Close and Allosteric Opening of the Polypeptide-Binding Site in a Human Hsp70 Chaperone BiP.” Structure 23(12): 2191–2203.

